# The glucocorticoid receptor as a master regulator of Müller cell gliosis in the diabetic retina

**DOI:** 10.1101/2023.09.06.556478

**Authors:** Anna M. Pfaller, Lew Kaplan, Madalena Carido, Felix Grassmann, Nundehui Díaz-Lezama, Farhad Ghaseminejad, Kirsten A. Wunderlich, Sarah Glänzer, Thomas Pannicke, Bernhard H.F. Weber, Susanne F. Koch, Boyan Bonev, Stefanie M. Hauck, Antje Grosche

## Abstract

Diabetic retinopathy (DR) is considered a primarily microvascular complication of diabetes. Müller glia cells are at the center of the retinal neurovascular unit and play a critical role in DR. We therefore investigated Müller cell-specific signaling pathways that are altered in DR to identify novel targets for gene therapy. Using a multi-omics approach on purified Müller cells from diabetic db/db mice, we found the mRNA and protein expression of the glucocorticoid receptor (GR) to be significantly decreased, while its target gene cluster was down-regulated. Further, oPOSSUM TF analysis and ATAC-sequencing identified the GR as a master regulator of Müller cell gliosis in DR. Cortisol not only increased GR phosphorylation. It also induced changes in the expression of known GR target genes in retinal explants. Finally, retinal functionality was improved by AAV-mediated overexpression of GR in Müller cells. Our study demonstrates an important role of the glial GR in DR and implies that therapeutic approaches targeting this signalling pathway should be aimed at increasing GR expression rather than the addition of more ligand.

**Graphical abstract:** 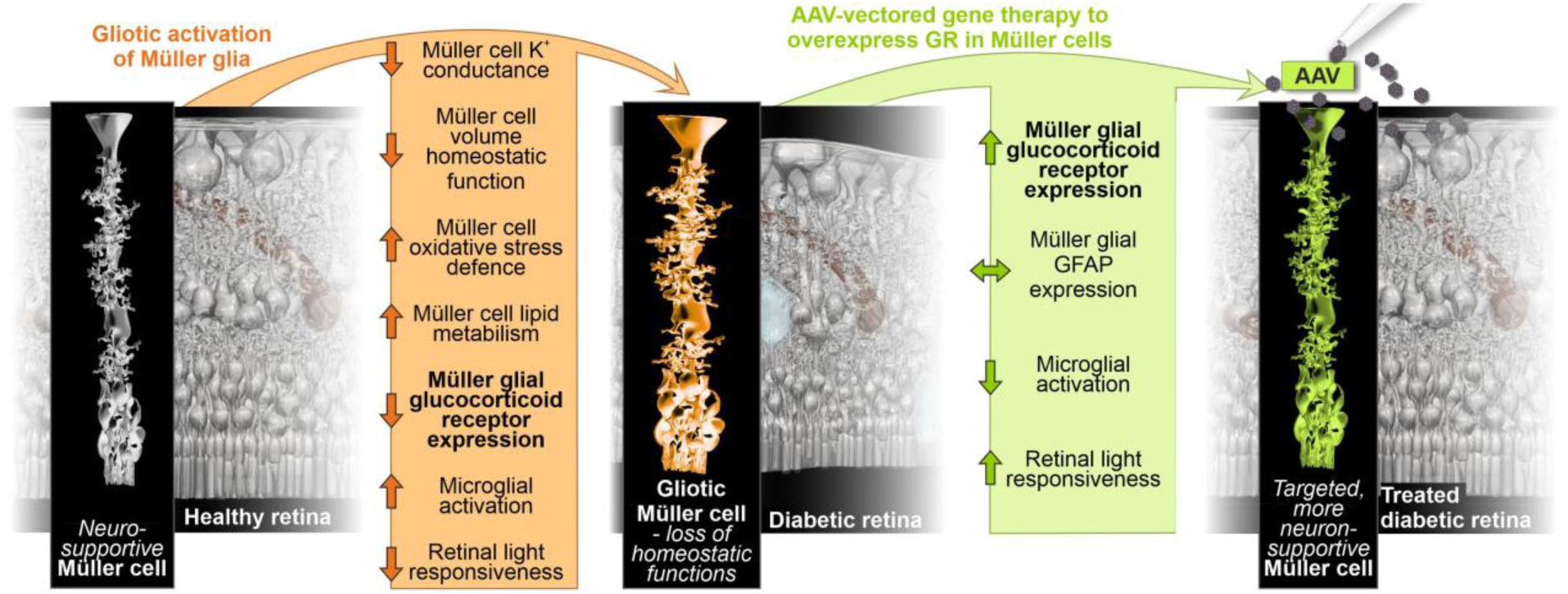

## Introduction

Diabetic retinopathy (DR), the leading cause of blindness in people of working age, is one of several complications of diabetes that affects retinal physiology and thereby visual acuity. Approximately one third of 246 million people with diabetes have evidence of DR, a third of whom develop severe retinopathy or macular edema (Cheung et al. 2010). DR can present as an early-stage non-proliferative form and can progress towards late-stage proliferative DR. Early DR is characterized by increased vascular permeability and capillary occlusion in the retinal vasculature. Microaneurysms, haemorrhages, cotton-wool spots and exudates may occur even in asymptomatic patients. The advanced stage of DR is dominated by neovascularisation, the outgrowth of new abnormal blood vessels. Patients experience severe visual impairment when the new abnormal vessels bleed into the retina and vitreous (Wang et al. 2018). Diabetic macular edema (DME), which can occur at any stage of DR, is the most common cause of vision loss in DR. A breakdown of the blood-retinal barrier (BRB) leads to intraretinal fluid accumulation and macular swelling (Wang et al. 2018). Changes in other retinal cell types leading to neurodegeneration even preceding microvascular remodeling are discussed (Sohn et al. 2016).

The pathology of DR depends on type and duration of the underlying diabetes, blood glucose levels and blood pressure (Tarr et al. 2013). DR-associated biochemical mechanisms are complex and affect cellular metabolism, signaling and growth factor release in retinal cell types. These pathways include an increased flux of glucose through the polyol and hexosamine pathways, accumulation of sorbitol and advanced glycation end products (AGEs), oxidative stress, activation of protein kinase C, chronic unfolded protein response activation, inflammatory responses, dysregulation of the renin-angiotensin system, and activation of the vascular endothelial growth factor (VEGF) signaling axis (Brownlee 2001, Cheung et al. 2010, Pitale et al. 2022). Accordingly, optimal control of blood glucose, blood pressure and lipids remains the basis for reducing the risk of DR development and progression (Cheung et al. 2010). In addition, current treatment strategies aim to target microvascular complications, and include intravitreal administration of pharmacological agents, laser therapy and vitrectomy. Intravitreal injection of anti-VEGF agents is currently the gold standard of therapy for early and advanced stages of DR (Wang et al. 2018). This therapy has been shown to improve visual acuity more effectively than laser treatment (Cheung et al. 2010). However, anti-VEGF therapy is also associated with limitations and adverse effects such as cataract formation, retinal detachment, vitreous haemorrhage, infection and possible loss of retinal neurons (Maharaj et al. 2008, Wirostko et al. 2008, Ford et al. 2011). In addition, efficacy depends on repeated (monthly) intravitreal injections because of the short half-life of the approved drugs increasing the likelihood of adverse complications such as endophthalmitis (Das et al. 2015, Dossarps et al. 2015). Finally, intraocular steroids are a traditional intervention for DR patients (Semeraro et al. 2019). As some patients become resistant to anti-VEGF therapy, these potent anti-inflammatory agents are becoming increasingly important (Wang et al. 2018). However, because of the relatively high incidence of adverse effects such as cataracts and glaucoma, corticosteroids are currently only a second-line option for patients who do not respond to other treatments (Wong et al. 2016). The implementation of gene therapy for DR, ideally as a one-time treatment, could be a long-term effective alternative to pharmacological strategies (Wang et al. 2020). However, respective approaches are still in the experimental stage by either targeting neovascularisation and vascular hyperpermeability or testing neuroprotective approaches in multiple DR-like animal models (Pechan et al. 2009, Haurigot et al. 2012, Zhang et al. 2015, Diaz-Lezama et al. 2016, Wang et al. 2020).

Although many of the cellular processes involved in DR have been extensively studied, the interplay between the different retinal cell types and the timing of these interactions are not fully understood (Wirostko et al. 2008). Currently, no animal model recapitulates the full multifactorial pathophysiology of DR. BKS.Cg-Dock7m+/+LeprdbJ (short db/db) mice develop type 2 diabetes (T2D) due to a mutation in the leptin receptor gene (Chen et al. 1996) and exhibit a number of abnormalities similar to those seen in human DR (Hammer et al. 2021). Elevated plasma insulin in homozygous mice occurs at 10 to 14 days of age (Coleman et al. 1974). They then become obese at 4 weeks and hyperglycemic at 4 to 8 weeks of age (Hummel et al. 1966). The first key features of DR, endothelial and pericyte loss as well as an altered neurovascular coupling, begin to develop in db/db mice at 10-12 weeks of age (Midena et al. 1989, Hanaguri et al. 2021). There is some disagreement as to when neuronal loss first occurs. Some studies describe a reduction in retinal layer thickness as early as 8 weeks of age (Tang et al. 2011, Bogdanov et al. 2014), while others noted key features of neurodegeneration, including a- and b-wave abnormalities in electroretinogram (ERG) recordings and TUNEL-positive cells in diabetic animals at 16 and 24 weeks of age (Bogdanov et al. 2014, Di et al. 2019). Only later, a breakdown of the BRB was reported and was clearly associated with increased apoptosis of neuronal cells, persistent glial activation and neovascularization (Cheung et al. 2005). Median survival of db/db mice is between 349 (male) and 487 (females) days (Sataranatarajan et al. 2016).

Müller cells are the major macroglia of the retina and are in contact with virtually all cell types of the retina, as well as blood vessels, the vitreous and the subretinal space (Bringmann et al. 2006, Bringmann et al. 2012, Reichenbach et al. 2013). They play a key role in maintaining retinal ion and volume homeostasis by mediating the transport of ions, water and various molecules (Bringmann et al. 2006, Wurm et al. 2011, Bringmann et al. 2013, Reichenbach et al. 2013). Together with microglial cells, Müller cells are involved in the immune response through various mechanisms, including their Toll-like receptors and secretion of cytokines and chemokines (Kumar et al. 2013), as well as by controlling retinal vascularisation and maintaining the blood-retinal barrier (Bringmann et al. 2006). Finally, Müller cells have neuroprotective properties through the release of neurotrophic factors, the uptake and recycling of glutamate or gamma-aminobutyric acid (GABA), and the secretion of the antioxidant glutathione (Bringmann et al. 2006, Bringmann et al. 2013), while also supporting the metabolism of retinal neurons (Reichenbach et al. 2013, Toft-Kehler et al. 2016, Ola et al. 2019, Calbiague Garcia et al. 2023). Accordingly, selective ablation of Müller cells has been shown to result in early photoreceptor degeneration, vascular abnormalities, breakdown of the BRB and neovascularisation in the retina (Shen et al. 2012).

Pathologic stress conditions such as diabetes lead to the induction of Müller cell gliosis. The extent to which gliotic activation of Müller cells affects neuronal survival is not fully understood. For example, it has been shown that glial glutamate transporter activity is decreased in Müller cells of diabetic rats (Li et al. 2002), leading to significantly increased intraretinal glutamate levels (Lieth et al. 2000). In addition, Müller cells from diabetic rats show a decreased POTASSIUM conductance of their plasma membrane, which is due to a redistribution of the Kir4.1 potassium channel (Pannicke et al. 2006). Consistent with this, POTASSIUM conductance was also decreased in Müller cells from patients with proliferative DR (Bringmann et al. 2002). Taken together, disruption of retinal POTASSIUM homeostasis together with dysfunction of glutamate transporters in Müller cells of diabetic patients may lead to neuronal hyperexcitation and glutamate excitotoxicity (Pannicke et al. 2006). Müller cells are a major source of VEGF in DR, and Müller cell-derived VEGF is a key driver of vascular lesion formation and hence inflammation in DR (Wang et al. 2010, Fu et al. 2015). Müller cells release also various cytokines and chemokines with angiogenic potential, including interleukin-6 (IL-6) (Yoshida et al. 2001, Rojas et al. 2010, Schmalen et al. 2021). Elevated IL-6 levels in diabetic patients have been associated with the development of ocular complications (Koskela et al. 2013). However, IL-6 has also been shown to prevent Müller cell dysfunction caused by hyperglycemia and to maintain proper neuronal function (Sappington et al. 2006, Yego et al. 2009).

Here, we report on the role of Müller cells in the context of DR, using the db/db mouse model of T2D. We identified pathways that are altered in Müller cells of the diabetic retina by analyzing changes in glial genome-wide chromatin accessibility in conjunction with the glial transcriptome and proteome during DR progression. This combinatorial approach identified the glucocorticoid receptor (GR) as a potential master regulator of glial changes in DR. Consequently, we performed AAV-mediated Müller cell-specific GR overexpression and demonstrated that the neuronal deficits observed in the db/db retina can be alleviated by this approach.

## Results

### Characterization of the retinal phenotype of db/db mice

To validate data from the literature and further characterize the retinal phenotype of the db/db mouse strain, morphometric analysis was performed on eyes from db/db and control animals at 3, 6, and 9 months of age (**Fig. 1A**). An important hallmark of diabetes is early changes in the integrity of the microvascular system, which are also observed in the progression of DR (Shin et al. 2014). Using trypsin-digested retinal flatmounts stained with hematoxylin and eosin (H&E), we analyzed the changes in the microvascular system of our db/db breed. Flat mounts from 3- and 6- month-old db/db animals and controls were used to quantify acellular capillaries (lagging pericytes and endothelial cells), but no significant differences were found at all ages analyzed (**Fig. S1**). A significant increase in the endothelial cell/pericyte ratio was calculated for 6-month-old diabetic animals compared to controls. In our db/db strain, only a mild decrease in pericyte number was observed in 6-month-old animals which became more pronounced in 9-month-old db/db mice (**Fig. 1B**). This age-dependent loss was not observed in wild-type controls at any age.

**Figure 1.**
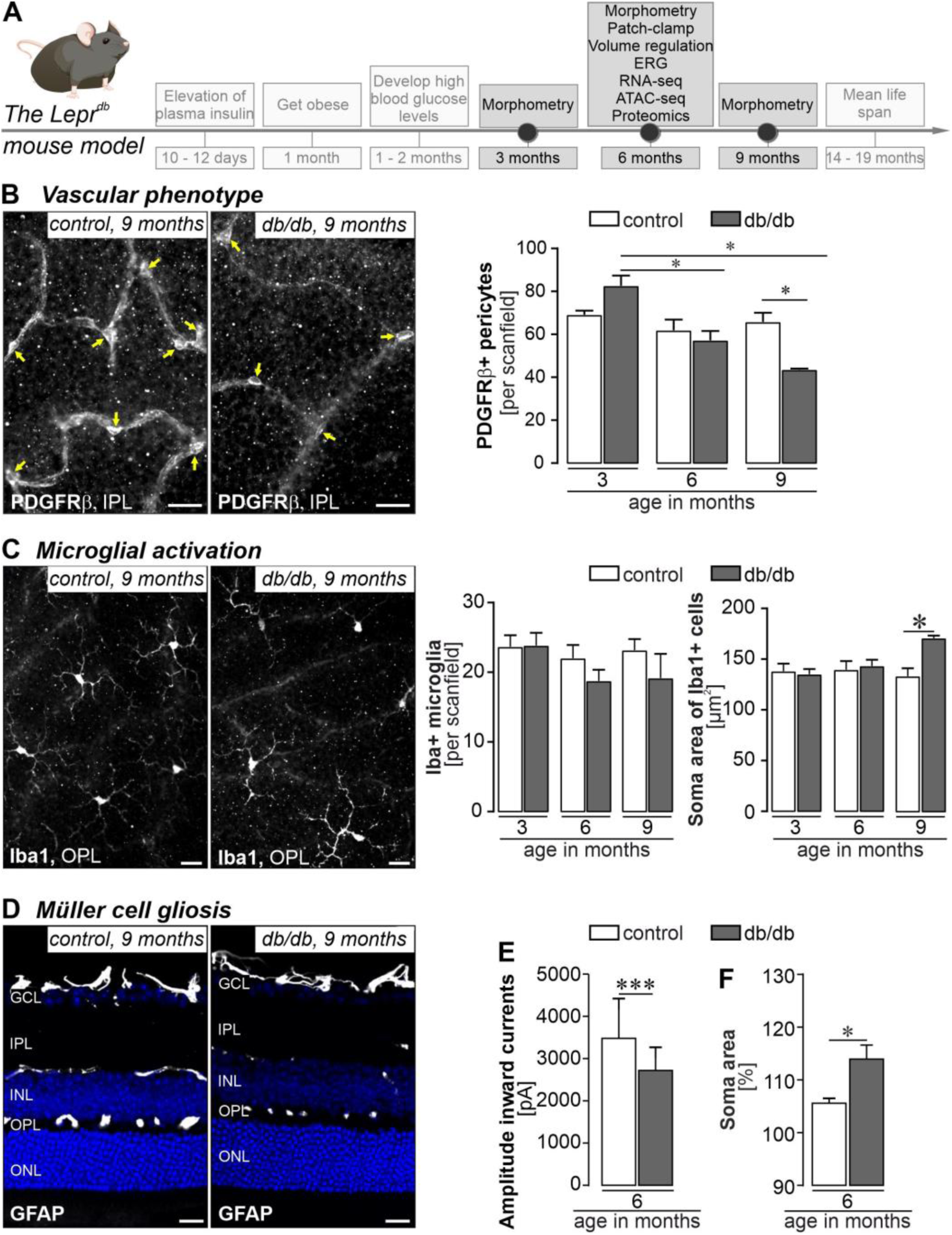
Characterization of the phenotype of the db/db (Lepr^db^) mice as an animal model for DR in T2D. *(A) Timeline of the development of features of T2D and DR in the db/db mouse model. Time points of data collection are highlighted as dots on the time line and respective readouts are listed in the boxes above*. *(B)* Left, *representative micrographs of PDGFRβ staining to delineate pericytes (yellow arrows) in retinal flatmount preparations of diabetic and control mice 9 months of age. Scale bar, 20 µm.* Right, *PDGFRβ-positive cells were counted per scan field (410 µm × 298 µm) of both genotypes. Bars represent mean ± SEM from n=3-6 animals per group. Unpaired t-test: *P<0.05*. *(C)* Left, *representative micrograph of Iba1 stainings of retinal flatmounts from 6-month-old diabetic and control mice*. *Scale bar, 20 µm.* Right, *Iba1-positive cells were quantified per scan field (410 µm × 298 µm). Z-scans through the whole thickness of the retina were performed and cells across all retinal layers were counted. In addition, the soma area of each microglia in a scan field was measured as an indicator of beginning microglial activation. Bars represent mean ± SEM and comprise data from 3-6 animals per age and genotype. Unpaired t-test: *P<0.05*. *(D) GFAP was only present in astrocytes residing in the nerve fiber layer, but not in Müller cells – neither in healthy controls nor in retinae from db/db mice 6-months of age*. *(E) Patch clamp recordings of isolated Müller cells from 6-month-old db/db mice demonstrated a significant reduction of amplitudes known to be primarily mediated by Kir4.1 potassium channels. Bars represent mean ± SEM and comprise data from ∼40 cells collected from 4 animals per genotype. Mann-Whitney-test: ***P<0.01*. *(F) The ability of Müller cells to compensate for hypoosmotic stress was tested in the retina of 6-month-old db/db mice*. *Vital retinal sections were exposed to hypoosmolar stress (60% osmolarity) for 4 minutes. Changes in Müller cell soma area, visualized by labeling with Mitotracker Orange, were measured as an indication of cell swelling. Bars represent mean ± SEM and comprise data from ∼15 cells collected from 2 animals per genotype. Mann-Whitney-test: *P<0.05*.

Next, we examined the number of Iba1-positive microglia/macrophages in retinae from db/db and control animals, but observed no significant differences between genotypes at any age examined (**Fig. 1C**). Still, the area of the microglial somata were significantly larger in the 9-months-old diabetic animals than in the age-matched control mice (**Fig. 1C**), suggesting enhanced microglial activation.

GFAP upregulation, indicating Müller cell gliosis, was not detected by immunostaining. Only astrocytes in the nerve fiber layer were immunopositive for GFAP (**Fig. 1D**). Patch clamp recordings from acutely isolated Müller cells from 6-month-old animals showed a significant decrease in potassium channel-mediated inward currents in db/db mice (**Fig. 1E**). Consistent with this, the ability of Müller cells in these diabetic animals to regulate volume was also severely diminished resulting in an increased soma size upon exposure to hypoosmotic stress (**Fig. 1F**). Thus, we confirm early functional changes in Müller cells at about the same age when the first signs of vascular changes appear, but before significant morphological changes in microglia were detected.

We then addressed the question whether the microvascular and glial alterations result in neurodegeneration. We quantified cell numbers in the ganglion cell layer (GCL), inner nuclear layer (INL), and outer nuclear layer (ONL) (**Fig. 2A**). We found no significant change in cell numbers in our db/db breed compared to control animals at any of the ages studied (**Fig. 2A**). To investigate neurodegeneration at the cell type level, we assessed the number of calretinin-positive cells representing ganglion cells and displaced amacrine cells in the GCL and amacrine cells in the INL (Lee et al. 2010, Lee et al. 2016), but again did not detect find significant differences between genotypes (**Fig. 2B**). Finally, we examined rod and cone photoreceptors known to be particularly susceptible to degenerative processes in multifactorial diseases such as DR (Kern et al. 2015). The total number of cones per scan field and the length of their outer segments were slightly reduced in retinae from diabetic animals at 9 months of age (**Fig. 2C**). We also measured the length of rod (outer and inner) segments based on PDE6B immunolabeling and found no difference between genotypes at any age investigated (**Fig. 2D**).

**Figure 2.**
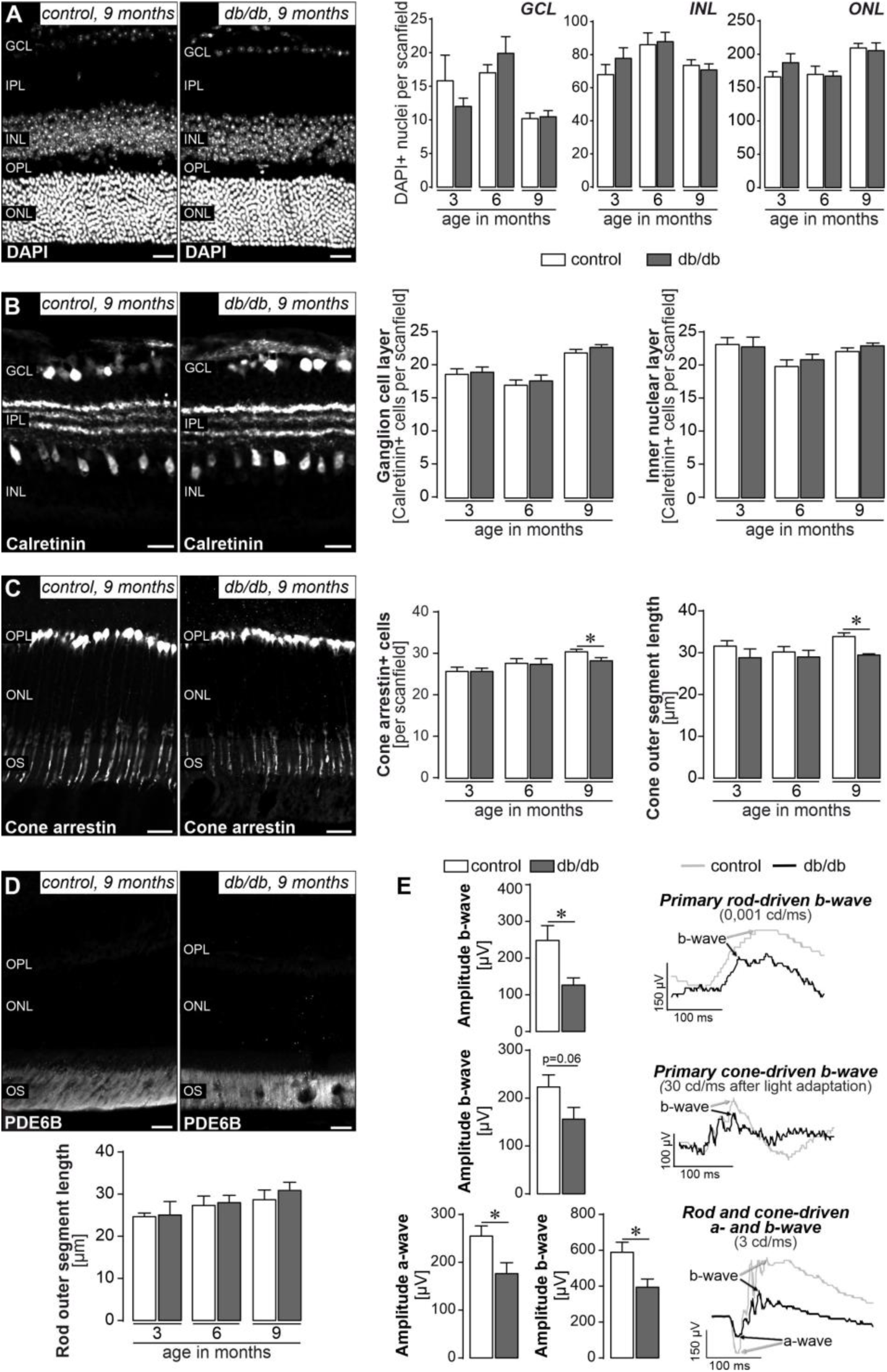
In-depth analysis of retinal degeneration in db/db mice. *(A)* Left, *representative micrograph of a DAPI-staining of the retina of 9-month-old diabetic and control mice.* Right, *quantification of DAPI-positive nuclei in the three nuclear layers of the retina does not reveal major cell loss even in 9-month-old db/db mice. Scan field: 68 µm × 200 µm*. *(B)* Left, *calretinin immunolabeling delineates ganglion and displaced amacrine cells in the ganglion cell layer (GCL) and amacrine cells in the inner nuclear layer (INL). Representative micrographs of retinae from 9-month-old diabetic and control mice are shown.* Right, *the number of calretinin-positive cells per scan field of the GCL and INL is plotted. Scan field: 200 µm × 200 µm*. *(C) Left, cone photoreceptors including their outer segments are visualized by a cone-arrestin staining for which representative results are presented from 9-month-old animals. Scale bar, 20 µm.* Right, *the number of cones and the length of their outer segments (OS) was assessed in retinae from both genotypes. Unpaired t-test: *P<0.05*. *(D)* Top, *representative micrograph of PDE6B immunoreactivity in rod OS in retinal sections from 9-month-old diabetic and control mice.* Bottom, *the rod OS length of retinae from db/db and control animals was measured*. *(A-D) Scale bars, 20 µm. Bars represent mean ± SEM and data from 3-4 animals per age and genotype. IPL, inner plexiform layer; OPL, outer plexiform layer; ONL, outer nuclear layer; OS, outer segments*. *(E) Electroretinogramm (ERG) recordings were performed on 6-month-old animals. Scotopic rod-specific b-wave, photopic cone-specific b-wave and mixed rod-cone–specific a- and b-wave were measured and quantified at 0,001 cd/ms, 30 cd/ms and 3 cd/ms, respectively. Right, representative ERG traces. Bars represent mean values ± SEM. 11 individual mice were measured per group. Unpaired t-test: *P<0.05*.

Next, we determined diabetes-associated changes on the functional integrity of the retinal tissue. Electroretinogram (ERG) recordings were performed in 6-months-old animals - a time point at which no major anatomical changes could yet be observed, but initial changes in the microvascular system (**Fig. 1B**) and Müller cell function (**Fig. 1D**) were detected. The rod (scotopic)-driven b-wave amplitude was significantly smaller in diabetic mice than in controls (**Fig. 2E**). The cone b-wave amplitude (photopic) was also reduced in diabetic mice, however, the effect was not statistically significant (p = 0.06). The mixed responses of cones and rods under mesopic light conditions were significantly reduced in diabetic mice for both a- and b-waves (**Fig. 2E**).

Overall, our data suggest that the progression of neurodegenerative processes as a hallmark of DR in the db/db mouse strain is slow, given that significant neurodegeneration occurs not before 9 months of age.

### Early changes in the glial transcriptome and proteome of diabetic retinae suggest metabolic dysregulation and altered growth factor signaling

The primary goal of this study was to define the gene expression signature of Müller cells in early DR. We isolated Müller cells and assessed Müller cell-specific responses for cross-comparison with microglia, vascular cells, and neurons from 6-month-old mice. Our data show the onset of vascular, glial, and even neuronal functional deficits but no significant morphological changes. RNA was extracted from all cell populations and subjected to RNA sequencing that was performed at 25-45 million reads per sample. The expression profiles of established marker genes were plotted and reveal successful enrichment of the respective cell populations (**Fig. S2A**). Interestingly, Itgam (also known as the microglial marker CD11b) was significantly upregulated in db/db mice. Principal component analysis (PCA) revealed that samples of the four cell types analyzed clustered separately based on their transcriptome (**Fig. S2B**), suggesting that cell identity dominates over diabetes-related changes in expression.

In light of their central role as a component of the retinal neuro-vascular unit primarily affected in DR, we focused our subsequent analysis on Müller cells. A PANTHER (released 20221013) overrepresentation test was used to perform pathway enrichment analysis on GO molecular functions of Müller cell-specific genes (i.e. 2-fold versus neuronal expression levels, p<0.05) that were significantly upregulated (117 genes) or downregulated (68 genes) in diabetic retina (**Tab. S1**). Genes related to aldehyde dehydrogenase (NAD+/NADP+), and glutathione transferase activity, in addition to several growth factor pathways, including brain-derived growth factor (BDNF), transforming growth factorβ (TGFβ) and insulin-like growth factor 1 (IGF1) as well as G-protein-mediated signaling, were upregulated in Müller cells from diabetic mice (**Figure 3A**, **Tabl. S1**). Pathways associated with downregulated genes included those involved with lipid metabolism (triglyceride ligase activity), extracellular matrix and cell-cell interactions (extracellular matrix binding, collagen binding, heparin binding, cadherin binding, actin binding), intracellular signaling (e.g. src homology (SH) regions 2 and 3 domain binding, phosphatidylinositol-3-kinase binding), and gene expression regulation (**Figure 3A**, **Tab. S1**).

**Figure 3.**
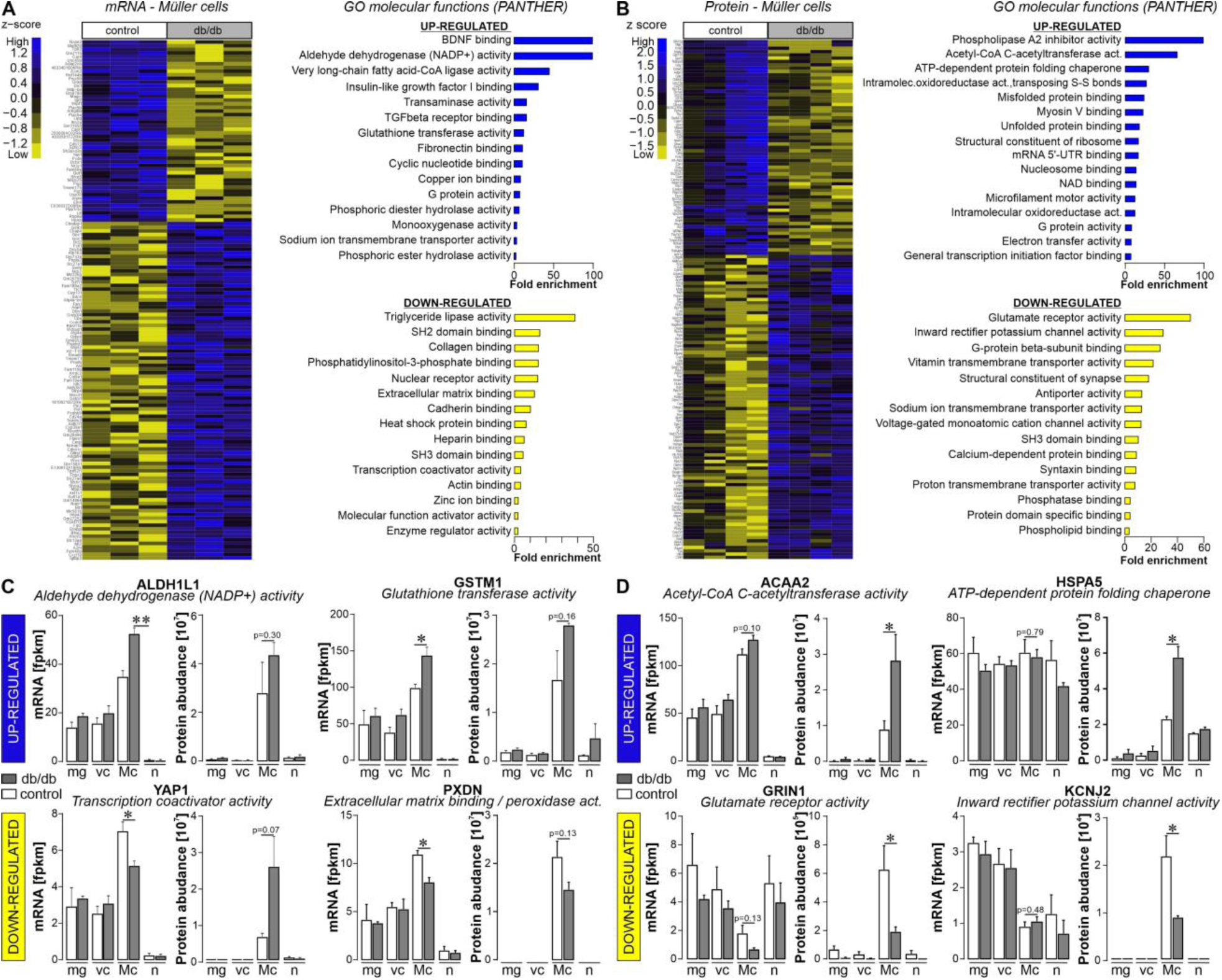
Expression landscapes at transcript and protein level of purified retinal Müller cells isolated from diabetic mice 6 months of age hint towards metabolic changes and altered growth factor signaling. *(A)* Left, *Differentially expressed genes of Müller cells isolated from 6-month-old db/db mice compared to controls (p<0.05; at least twofold difference), which additionally showed Müller cell specificity (p<0.05 compared to expression in neurons; at least twofold difference is plotted in the heatmap). The heatmap includes three biological replicates per genotype to illustrate the degree of heterogeneity in gene expression within each genotype.* Right, *GO molecular functions (based on fold enrichment, at least twofold) of significantly up- or down-regulated genes in Müller cells from 6-month-old db/db mice as determined by pathway enrichment analysis using PANTHER*. *(B)* Left, *filtering (P< 0.05; at least twofold difference) was performed to identify proteins differentially regulated in Müller cells from 6-month-old diabetic mice.* Right, *molecular functions driven by proteins significantly down- or up- regulated in Müller cells from 6-month-old db/db mice determined by pathway enrichment analysis using PANTHER*. *(C) Transcript and protein expression of select candidate genes central to the pathways as identified from differentially expressed Müller cell-specific transcripts shown in (A) are plotted across all cell types to illustrate Müller cell-specificity of the diabetes associated gene expression. Bars represent the mean ± SEM of three biological replicates for RNA-seq data and 4 biological replicates for protein abundances, respectively. Unpaired t-test: *P<0.05, **P<0.01*. *(D) Transcript and protein expression of select candidate genes central to the pathways as identified on the proteome profiles shown in (B) are plotted across all cell types to illustrate Müller cell-specificity of the diabetes associated gene expression change. Bars represent the mean ± SEM of three biological replicates for RNA-seq data and 4 biological replicates. Unpaired t-test: *P<0.05*. *(C, D) mg, microglia; vc, vascular cells; Mc, Müller cells; n, neurons*.

We also subjected retinal cell populations purified by magnetic-activated cell sorting (MACS) to unbiased label-free liquid chromatography-mass spectrometry proteome profiling. To maximize comparability, the same neuronal, microglial, vascular and Müller cell marker genes were selected as previously used to validate cell enrichment in the RNA-seq dataset. This convincingly confirmed the successful separation of the different retinal cell populations (**Fig. S2C**). PCA based on the cell type-specific proteomes showed that cell populations formed four clearly separated clusters (**Fig. S2D**). The proteins which were significantly up (92 proteins)- or down (66 proteins)-regulated in Müller cells of 6-month-old diabetic mice were identified (**Fig. 3B**, **Tab. S2**). Analysis of the enrichment of down-regulated proteins by PANTHER shows down-regulation of adenosine triphosphatase (ATP) and transporter activity (**Fig. 3B**). In contrast, pathways related to RNA, protein binding, ligase, and transferase activity were associated with proteins upregulated in Müller cells from diabetic mice (**Fig. 3B**).

Since we generated the transcriptomic and proteomic data in a highly comparable manner with respect to the cell isolation protocols and the mouse strain used, we asked to what extent changes in gene expression at the transcriptional level are actually reflected in the proteomes of the cells. A total of 883 Müller cell-specific genes (56.4%) present in both data sets showed a congruent expression profile at the mRNA and protein level (**Fig. S3A**). Focusing on the genes that were differentially expressed in Müller cells of the diabetic retina resulted in similar proportion of genes (57.3%) that were found to have a consistent regulation at the mRNA and protein level (43 out of 75 differentially expressed genes (DEGs, **Fig. S3B**). This partial mRNA/protein discrepancy for some DEGs is also reflected in the expression profiles of selected candidates that were central to pathways discussed above (**Fig. 3C,D**). While upregulation of aldehyde dehydrogenase 1 family member L1 (*Aldh1l1*) or glutathione S-transferase mu 1 (*Gstm1*) was specific to Müller cells and consistent in both transcript and protein expression, significant downregulation of Yes1-associated transcriptional regulator (*Yap1*) at the RNA level was not reflected in protein expression, but YAP1 protein levels were higher in db/db Müller cells compared to wild-type cells. Finally, the down-regulation of peroxidasin (*Pxdn*), a heme-containing peroxidase that is secreted into the extracellular matrix, was consistent again for transcript and protein expression. Focusing on the DEGs driving the pathway analysis shown in **Figure 3B**, we found a similar discrepancy between transcript and protein expression. Consistent expression was found for mitochondrial acyl-CoA hydrolase (*Acaa2*), an enzyme that catalyzes the final step of mitochondrial fatty acid beta-oxidation, which was significantly and specifically upregulated in Müller cells (**Fig. 3D**). Also for glutamate ionotropic receptor NMDA Type Subunit 1 (*Grin1*), a significantly down-regulated protein in Müller cells, the regulation of the transcript matches the protein. While *Grin1* transcripts were rather equally detected in all cell populations, the protein was almost exclusively present in Müller cells. In contrast, transcripts of heat shock 70kDa protein 5 (glucose-regulated protein, 78kDa; *Hspa5*) and inward rectifier potassium channel 2 (alias Kir2.1, *Kcnj10*) were not differentially expressed or even specific to Müller cells, while the respective protein was significantly up- or down-regulated and specific to the Müller cell population when compared to microglia, vascular cells or neurons (**Fig. 3D**). The cytoskeletal-associated protein 4 (*Ckap4*), a novel RNA-binding protein (Lyu et al. 2019), is the only Müller cell-specific gene we found to be consistently and significantly dysregulated at both mRNA and protein level (**Fig. S3B, C**, **Tabs. S1, S2**).

### The glucocorticoid receptor was identified as factor potentially driving changes in the Müller cell gene expression pattern in the diabetic retina

To identify key regulators of Müller glial changes in DR progression, we investigated changes in DNA accessibility by performing ATAC-seq on purified Müller cells from 6-month-old mice and subjected these data to transcription factor (TF) binding motif analysis. Motifs for the glucocorticoid receptor (gene ID: Nr3c1) stand out as being more accessible in Müller cells isolated from diabetic retinae compared to controls (**Fig. 4A**). In contrast, motifs for TFs that are known to determine and maintain photoreceptor identity (*Crx, Otx1, Otx2*) were less open in cells isolated from db/db retinae (Ghinia Tegla et al. 2020). Since some contamination of our Müller cell population by photoreceptors cannot be avoided (Grosche et al. 2016), this latter finding may also reflect early changes in photoreceptor gene expression signatures long before functional (**Fig. 2E**) or even morphological (**Fig. 2C**) findings are observed. Finally, motifs for two members of the NFI family - *Nfix* (var. 2) and *Nfib,* as well as *Ctcf* were enriched in regions which were accessible in both control and diabetic cells (**Fig. 4A**).

**Figure 4.**
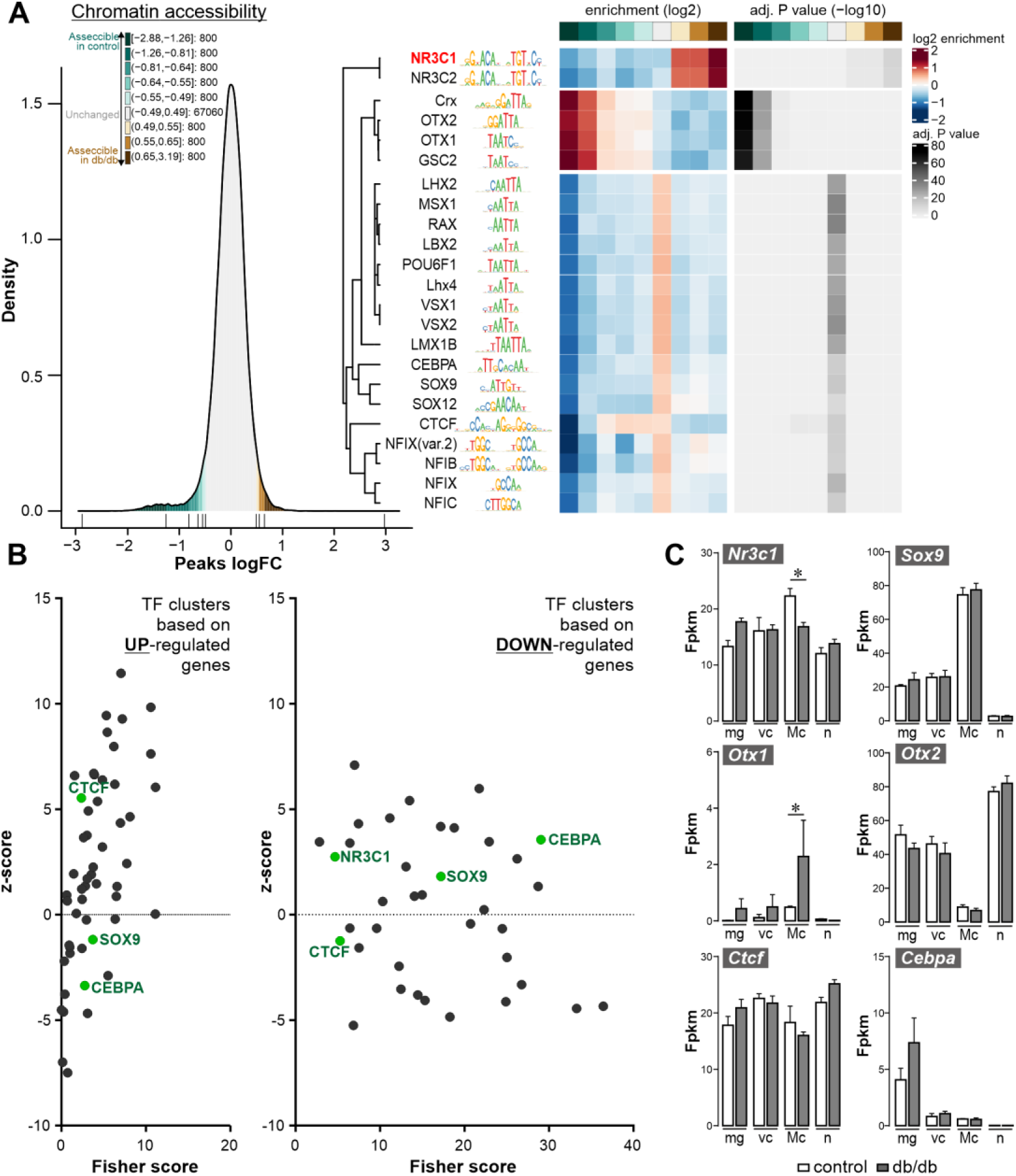
Identification of the glucocorticoid receptor (gene ID: Nr3c1) as a putative key regulator of glial transcriptomic changes in diabetic mice on basis of chromatin accessibility and RNA expression profiles. *(A) Chromatin accessibility was assessed by ATAC sequencing on purified Müller cells from 6-month-old mice of both genotypes. Data were subjected to transcription factor binding motif analysis.* Left: *Density plot showing the distribution and color code of bins comprising an equal number of peaks corresponding to fold changes in chromatin accessibility between db/db and control mice. Numbers represent the log2 fold change range (db/db vs control) and the corresponding number of peaks per bin.* Right: *cluster analysis and heat map of transcription factor (TF) binding motifs significantly enriched when comparing chromatin accessibility of Müller cells from db/db mice with their counterparts from wild-type animals. Bins in shades of green indicate decreased accessibility, while shades of brown indicate increased accessibility. Motifs enriched in the gray bin are those with unchanged chromatin density between genotypes*. *(B) TF clusters possibly involved in regulatory changes associated with DR progression as identified by oPOSSUM-3 analysis. Genes/transcripts identified by RNA seq to be significantly up- or down-regulated in Müller cells of 6- months-old diabetic mice were submitted to oPOSSUM-3 analysis (Ho Sui et al. 2005). Only TFs that were expressed at transcript level in Müller cells are included. TFs identified in both ATAC-seq and oPPOSSUM-3 analysis are highlighted in green*. *(C) Expression levels of TFs highlighted in green in (B) and of OTX1 and OTX2 as two examples of the TFs that were prominent in (A) by a high-level reduction in the accessibility of their DNA-binding domains. Data are from the RNA-seq dataset performed on MACS-sorted cells (**see*** Fig. 3*). Bars represent mean ± SEM. Dots represent biological replicates. Unpaired t-test: *P<0.05. Mg, microglia; vc, vascular cells; Mc, Müller cells; n, neurons*.

Next, the search for key regulators of Müller glial changes in DR progression was approached from the mRNA perspective. An oPOSSUM-3 TF binding site cluster analysis (Kwon et al. 2012) based on differentially regulated genes in Müller cells from 6-month-old diabetic mice as identified by the RNA-seq experiment identified multiple gene clusters that are upregulated in Müller cells, e.g. *Sp1*, *Ebf1*, *Egr1*, *Zfx*, *Sox9* and *Foxd1* (**Fig. 4B**, **Tab. S1**), and gene clusters related to down-regulated transcripts including *Sox9*, *Rora*, *Foxf2*, *Srf* and *Nr3c1* (**Fig. 4B**, **Tab. S1**).

Of note, several TFs, including *Nr3c1, Sox9, Ctcf,* and *Cebpa* (highlighted in green, **Fig. 4**), were identified via oPOSSUM and ATAC-seq approaches, making them potentially interesting candidates as master regulators of glial gene expression signatures in DR. *Nr3c1* (GR) transcripts, while present across all cell types, were most abundant in Müller cells, and were significantly reduced in 6-month-old db/db mice as compared to age-matched controls (**Fig. 4C**). *Sox9* transcripts were even stronger enriched in Müller cells than that of the GR, but no change of its expression level was obvious between genotypes investigated. *Ctcf*, in fact a putative target gene of *Nr3c1*, was also uniformly expressed in all retinal cell types, with the least of its transcripts detected in Müller cells. In Müller cells from diabetic mice, *Ctcf* expression was slightly lower than in controls, although this trend did not reach significance. *Cebpa* was found to be expressed specifically in microglia with a trend of upregulation in cells isolated from 6 months old mice. Finally, two out of the four TFs, for which highly significant closure of DNA binding motifs was identified (**Fig. 4B**), were also added. *Otx1* transcripts were expressed at low levels in Müller cells (**Fig. 4C**). Interestingly, significantly more transcripts were detected in cells isolated from 6-month-old diabetic mice. In contrast, Otx2, which together with the well-established photoreceptor-specific TF *Crx*, is a known driver of photoreceptor differentiation and controls the expression of rhodopsin in rods (Kaufman et al. 2021, Langouet et al. 2022), was significantly enriched in the neuronal population (Fig. 4C). However, no significant effect of the diabetic condition on its expression level was detected in any of the cell types, including the rod-rich neuronal population (Grosche et al. 2016).

Ultimately, the GR stands out from all TFs mentioned because (i) its DNA binding motifs show altered accessibility with progression of DR, (ii) it could also be identified as a potential master regulator via our Müller cell-specific RNA-seq dataset, as target genes of the GR were enriched among the differentially expressed Müller cell-specific genes, and (iii) the GR is expressed at comparatively high levels in Müller cells.

### In depth characterization of Nr3c1 as potential master regulator of Müller cell gene expression

Given the results from ATAC-seq and oPOSSUM analysis, we first aimed to confirm that the glucocorticoid receptor (GR; gene ID: Nr3c1) is expressed in Müller cells and is differentially expressed in the diabetic retina. GR-immunoreactivity was clearly localized to Müller cell somata in the inner nuclear layer and, the staining intensity seems weaker in sections from 6-month-old diabetic mice (**Fig. 5A**). Super resolution imaging by STED shows little overlap of GR signals in Müller cell somata with signals from cytoplasmic glutamine synthetase, but a broad overlap with nuclei visualized by the DAPI staining. Interestingly, GR does not seem to be present in nucleoli (**Fig. 5A**), but locates primarily in the less condensed euchromatin of the nucleus consistent with the function of a transcription factor. Western blots performed on purified retinal cell populations from control and diabetic mice at 6-months of age confirmed the findings from the immunostainings that the GR is also down-regulated at protein level in Müller cells of the diabetic retina (**Fig. 5B**, **Fig. S4 A**). Interestingly, we detected a significantly higher plasma cortisol level in 6-month-old animals of our breed by ELISA (**Fig. 5C**). Next, we asked whether the remaining GR could be more active due to a surplus of available ligand. To this end, we detected enhanced GR-phosphorylation in whole retina protein extracts from 6-month-old animals and found a trend of increased activation (**Fig. 5D**, **Fig. S4 B**).

**Figure 5.**
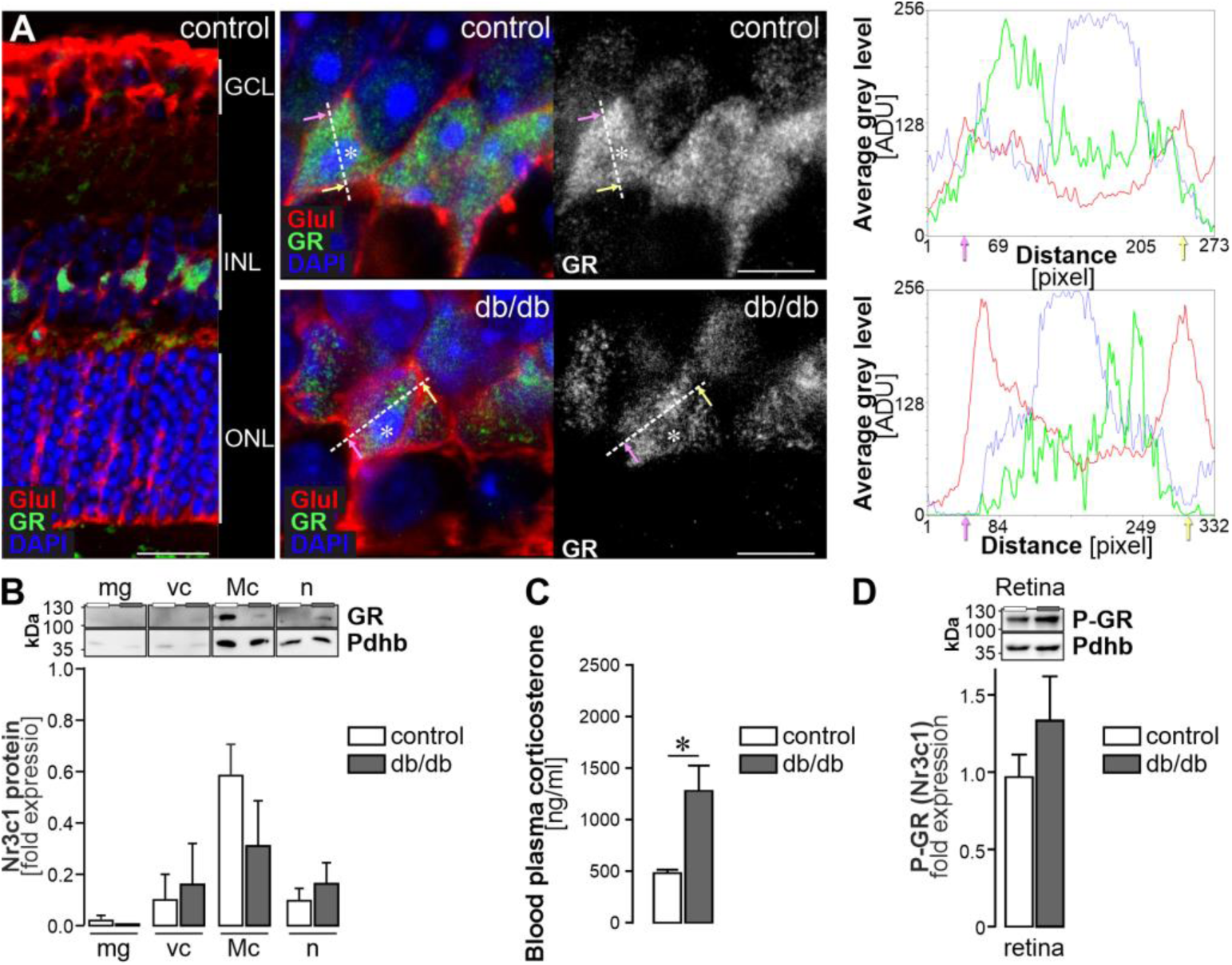
The glucocorticoid receptor (gene ID: Nr3c1) is specifically expressed in Müller cells and modulates glial gene expression. *(A)* Left, *confocal image of a GR labeling in retinal section of a 6 -month-old mouse. Scale bar, 20 µm*. Middle, *STED images of a GR staining in the inner nuclear layer (INL). Müller cells were co -stained for glutamine synthetase (GLUL). Scale bars, 5 µm.* Right, *line plots of mean grey values for each channel (red – GLUL as Müller cell marker residing in the cytoplasm; green – GR; blue – DAPI highlighting the DNA condensed in the nucleus). The dashed line indicates the plane of the line scan in the respective micrographs. Pink and yellow arrows indicate the orientation of data plotted in the histograms in relation to the actual line set in the micrograph. Asterisks highlight nucleoli. GCL, ganglion cell layer; INL, inner nuclear layer; ONL, outer nuclear layer*. *(B) Western blot to detect GR in purified retinal cell types isolated from 6 -month-old mice. As the protein yield per cell population isolated per two pooled retinae is very low, the whole protein extract per cell pellet was loaded. To enable quantification of the GR band, the signals were normalized to the house keeper PDHB as done in previous studies implementing MACS-purified retinal cell types (Pauly et al, 2019). Bars represent mean ± SEM (n=6)*. *(C) The corticosterone level was measured in the blood plasma of diabetic and control animals at an age of 6 months via ELISA. Results of n= 4 to 5 animals per group are plotted as mean ± SEM. Unpaired t-test: *P<0.05*. *(D) Western blot of phosphorylated GR performed on whole retinal extracts from animals 6 months of age*. *Bars represent mean ± SEM (n=4 animals per genotype)*.

To determine whether increased exposure to GR ligands affects the expression profile of Müller cells, retinal explants of wild-type mice were kept in culture and cortisol was supplemented (**Fig. 6A**), which led to higher levels of GR activation indicated in the its enhanced phosphorylation (**Fig. 6B**, **Fig. S4 C**). For unbiased analysis of the putative GR gene regulatory network, we performed mass spectrometric protein profiling on explants after 2 days of cortisol treatment. 72 proteins were found to be significantly upregulated and 61 were down-regulated (**Fig. 6C**). Pathway enrichment analysis revealed genes involved in ion and vesicular transport were down, while oxidative stress defense and regulation of inflammatory pathways were up-regulated upon cortisol treatment (**Fig. 6D**). Eleven of the upregulated genes were potential direct targets of GR as reported by the three databases JASPER, ENCODE, and CHEA (Davis et al. 2018; Mathelier et al. 2014; Rouillard et al. 2016) and matched with gene expression changes upon cortisol treatment. Cross-validating the Müller cell-specificity of these 11 candidates, we checked published scRNA-seq resource data from mouse retina (Macosko et al. 2015) (**Fig. 6E**).

**Figure 6.**
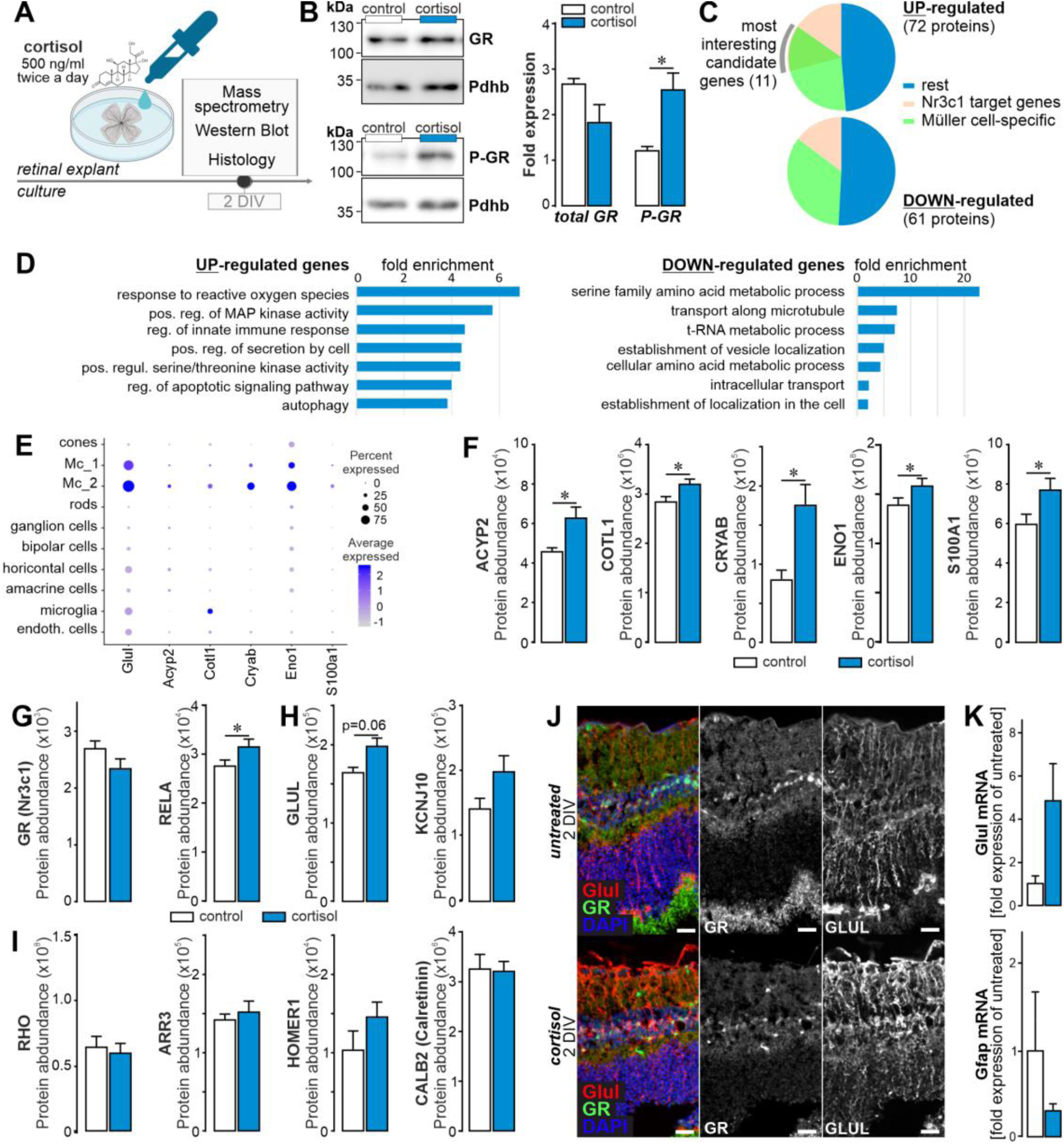
Cortisol treatment of wild-type mouse retinal explants reinforces GR signaling and enhances glial homeostatic gene expression. *(A) Experimental scheme of retinal explant cultures with persistent GR activation*. *(B)* Left: *Representative Western blots performed on whole retinal explant protein extracts when cortisol (500 ng/ml) was added twice daily.* Right: *Quantification of the total amount of GR and P-GR protein levels at 2 days of cortisol treatment. PDHB served as housekeeper to which the GR a nd P-GR were normalized to. Results of n=3 explants per group are plotted as mean ± SEM. Unpaired t-test: *P<0.05*. *(C) Retinae were cut into half. One half remained untreated and was kept under standard culture conditions. The other half was treated with cortisol (500 ng/ml, supplemented twice a day). Mass spectrometric profiling of explants after cortisol treatment over 48 h revealed 133 differentially expressed proteins. Using JASPER, ENCODE and CHEA databases, we identified those proteins whose genes are putative targets of GR and checked if those were then Müller cell -specific basing on our own RNA-seq data of purified retinal cell types. Eleven overlapping proteins were found among the up - regulated candidates, but none among the down-regulated upon cortisol treatment*. *(D) Pathway enrichment analysis of differentially expressed proteins from (D) using STRING*. *(E) Reanalysis of single cell RNA-seq data from Macosko et al. (Macosko et al. 2015) confirmed Müller cell-specific expression of many of the 11 GR target genes found to be up-regulated upon cortisol treatment*. *(F) Protein quantification via mass spectrometry (n=6) of putative Müller cell -specific GR target genes as identified by filtering of data in (D). Unpaired t-test: *P<0.05*. *(G, H, I) Protein expression levels of select candidates are plotted on basis of quantitative mass spectrometric data collected from cortisol-treated retinal explants. Besides Nr3c1, its interaction partner RELA was chosen (H), and additionally Müller cell (I) and neuronal marker genes (J). GLUL, glutamine synthetase; KCNJ10, Kir4.1; RHO, rhodopsin; ARR3, cone arrestin; HOMER1, homer scaffold protein 1; CALB2; calretinin*. *(J) Sections from retinal explants were stained for the Müller cell marker glutamine synthetase (GLUL) and GR*. *(K) Quantitative real-time PCR (qPCR) of Müller cells purified form retinal explant cultures at 2 DIV was performed to determine mRNA levels of Glul and Gfap. Bars represent mean ± SEM (n=3 for each condition). Scale bars, 20 µm*.

Five candidate genes were predominantly expressed in Müller cells and were significantly up-regulated upon cortisol treatment (**Fig. 6F**). This supports the idea that modulation of GR signaling can directly affect Müller cell-specific genes. Finally, we analyzed GR protein and interacting *Rela* expression in the cortisol-treated retinal explants. GR/Nr3c1 expression was slightly reduced (confirming the assumption of an autoregulatory repression), while that of Rela seemed to be increased (**Fig. 6G**). Confirming earlier reports, Müller cell genes such as glutamine synthetase and Kir4.1 were expressed at higher levels upon cortisol treatment (**Fig. 6H**) (Toops et al. 2012). None or moderate changes were observed for marker genes of rods (rhodopsin), cones (arrestin3), synapses (HOMER1) or ganglion and amacrine cells (calretinin) (**Fig. 6I**). Accordingly, cortisol treatment may help to maintain Müller cells in their neuron-supportive state preventing gliosis induction, which is partially reflected by the stable or even slightly improved expression of neuronal markers (e.g. HOMER1). Immunostaining of explants 2 days *in vitro* (DIV) support that GLUL expression seems more stable in cortisol treated samples (**Fig. 6J**). To confirm this finding, qPCR on purified Müller cells from retinal explant cultures at 2 DIV was performed and confirmed that *Glul* expression was higher in cortisol treated samples than untreated controls (**Fig. 6K**).

Having demonstrated Müller cell-specific expression and down-regulation of *Nr3c1* in early DR and the finding that enhanced GR-mediated signaling promotes a potentially more neuron-supportive Müller cell phenotype in cortisol-treated retinal explants, we revisited our ATAC- and RNA-seq data to check whether the regulation pattern of known GR target genes is in line with our assumption that reduced GR activity is a key driver of Müller cell gliotic alterations. Confirming the data presented in **figure 4**, we found an enhanced accessibility at peaks associated with *Nr3c1* motifs in 6-months old diabetic animals (**Fig. 7A**). Next, we analyzed whether the altered DNA accessibility at Nr3c1 binding motifs is reflected by changes in mRNA levels of known GR target genes (**Fig. 7B**). The data of GR target gene expression and their regulation in Müller cells of diabetic retinae was rather heterogeneous. While a number of target genes (e.g., Abhd2, Ptch1, Clrn1, Car1, Sfxn5) are downregulated, suggesting that GR may act as a transcriptional activator, we also found that GR target genes, e.g., Acyp2, Cryab, Aldh1l1, Fstl3, that were up-regulated. This suggests that GR acts as an activator or repressor depending on the DNA landscape and potential interaction with additional transcription factors on these target genes in Müller cells.

**Figure 7.**
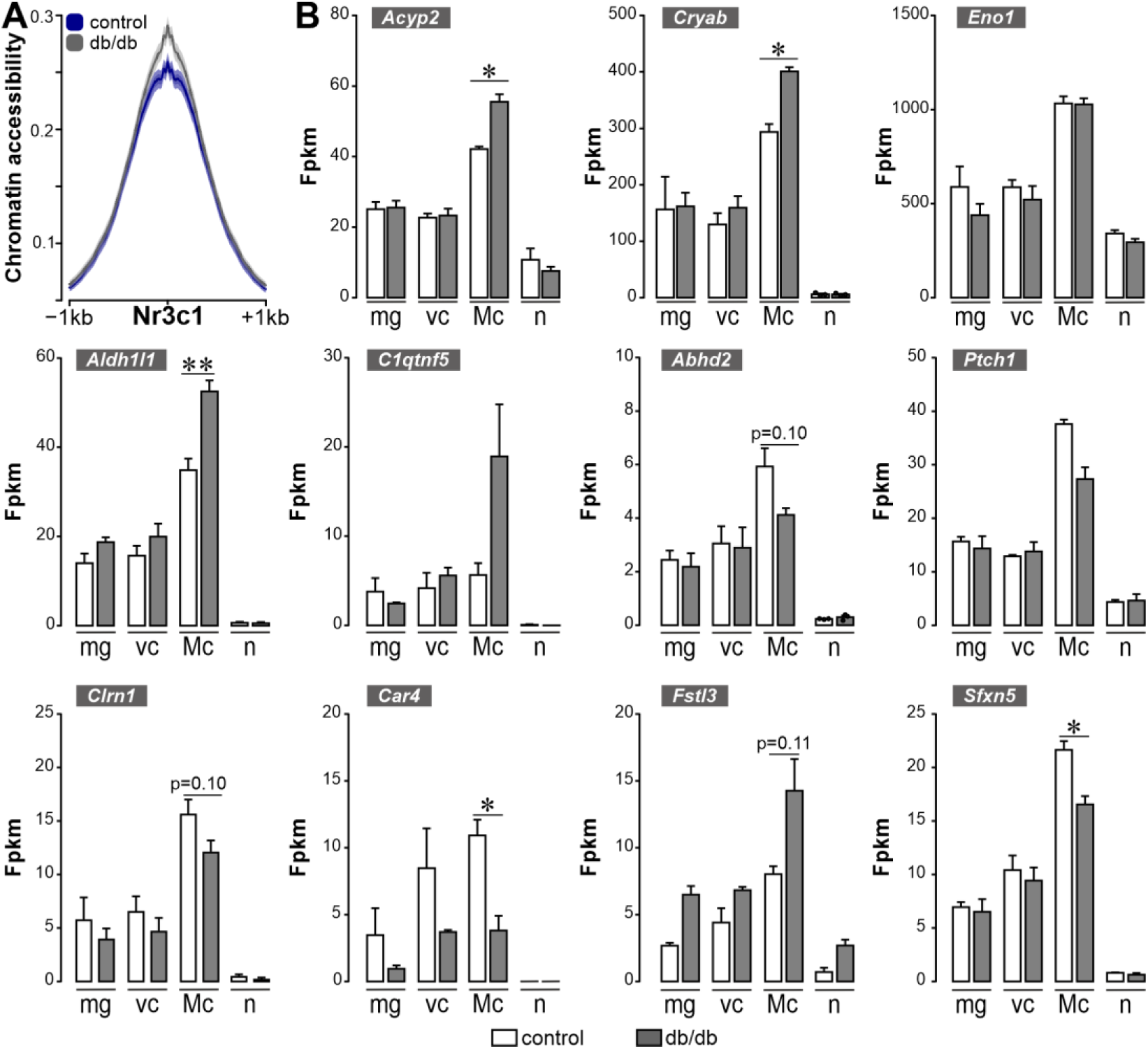
Regulation pattern of GR target genes in Müller cells of the diabetic mouse. (A) Accessibility at peaks associated with the Nr3c1 motif is plotted comparing results from Müller cells isolated from 6-month-old control and db/db animals. (B) Plots of transcript expression of GR target genes as determined by the RNA-seq experiment on purified retinal cell types from 6-month-old animals. Bars represent mean ± SEM. Unpaired t-test: *P<0.05;**P<0.01.

### Therapeutic intervention by overexpression of Müller cell-specific glucocorticoid receptor in db/db mice

In the final set of experiments, we attempted to restore high levels of GR expression in Müller cells of the diabetic retina. To achieve this, we implemented AAV9-vectorized GR overexpression driven by the GFAP promotor (**Fig. 8A**). Since GFAP is upregulated in Müller cells of the diabetic retina (**Fig. S3**), the GR overexpression construct should be active primarily in diseased tissue and, importantly, the GFAP promoter should not be targeted by the autoregulatory loop that down-regulates GR expression in the presence of high levels of corticosterone, as happens with endogenous GR. Immunostaining for the reporter EGFP in eyes one month after intravitreal injection of either AAV or PBS (sham control) confirmed the glial specificity of the construct (**Fig. 8B**). Importantly, a moderate up-regulation of GR was observed in GLUL-coexpressing Müller cell bodies in the INL (**Fig. 8B**). Note that no change in GR expression was observed in the nerve fibre/ganglion cell layer where astrocytes, potentially also targeted by the GFAP promoter-driven construct, reside (**Fig. 8B**). This suggests that the effects of the treatment are mainly due to GR overexpression in Müller cells.

**Figure 8.**
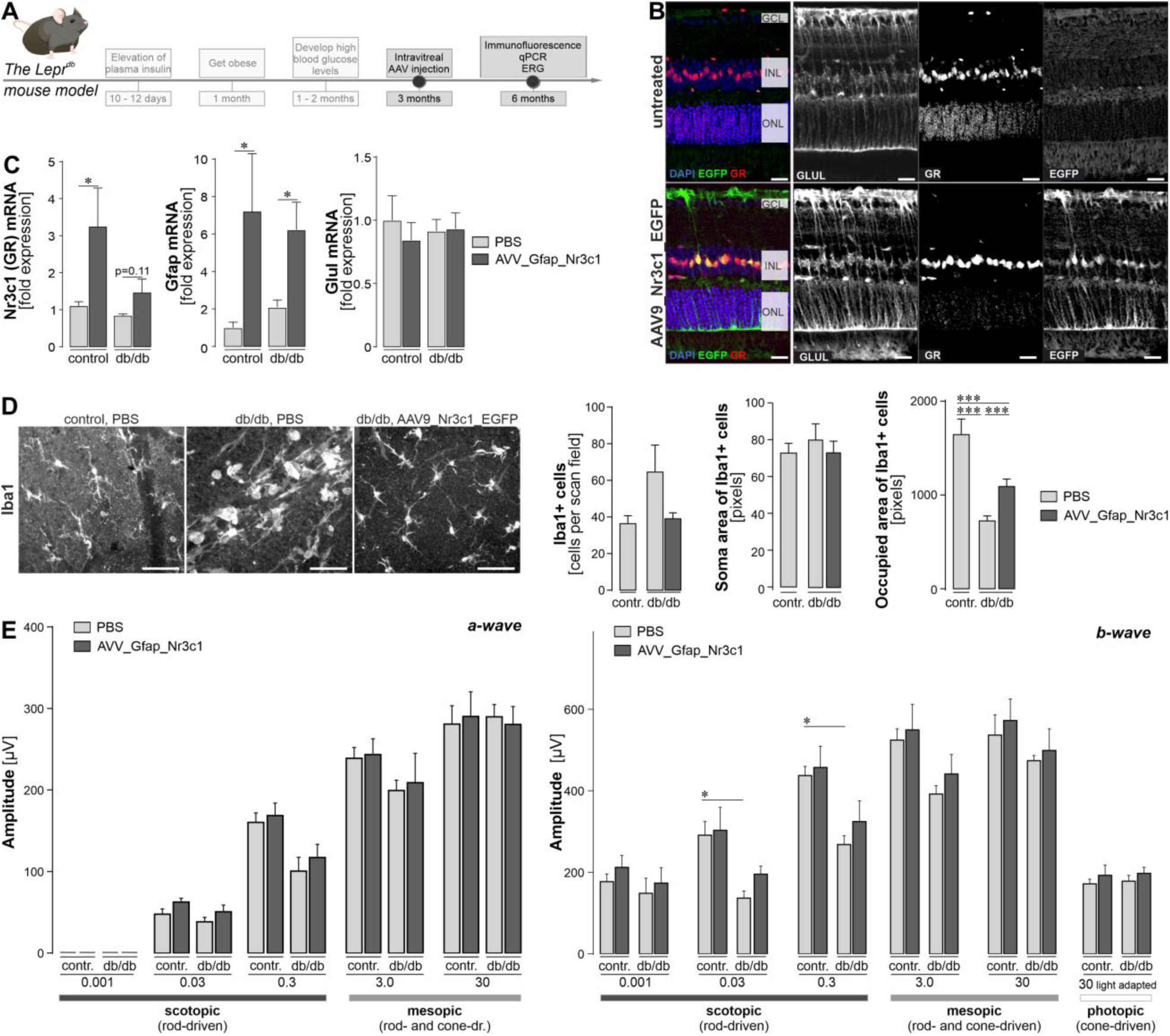
Overexpressing GR (gene ID: Nr3c1) in Müller cells of the diabetic mouse retina in vivo. (A) Experimental design and the time line of treatment and readouts in relation to the development of diabetes in the db/db mice. (B) 2.5 × 10^13^ GC/ml AAV9-Nr3c1-EGFP particles were injected intravitreally in 3-month-old diabetic and control mice. Tissue was collected 3 months thereafter and stained for the Müller cell marker glutamine synthetase (GLUL). EGFP indicated successful viral transduction. GCL, ganglion cell layer; INL, inner nuclear layer; ONL, outer nuclear layer. Scale bar, 20 µm. (C) qPCR on MACS-sorted retinal cells isolated from 6-month-old mice, 3 months after single AAV injection, was performed. Only Müller cell-specific gene expression is shown. Bars represent mean values ± SEM. n = 6-7 in controls and n = 4-5 for db/db mice. Unpaired t-test: *P<0.05. (D) Assessment of microglial responses upon AAV-treatment reveals reduced microglial activation in retinae with Müller glia-specific GR overexpression. Left, representative images of micrographs depicting the Iba1-staining in retinal flatmounts that were used to assess microglial morphological alterations. Right, quantification of microglial cell numbers, soma size and the area occupied by their processes. The bigger the soma area and the smaller the occupied area, the more activated the microglia are. Ordinary one-way ANOVA with Dunnett’s multiple comparisons test, ***P<0.001. (E) Electroretinogram recordings were performed 3 months after AAV injection. The contralateral eye received an equal volume of PBS as sham control Primarily rod (scotopic)- or cone-driven responses (photopic) or mixed responses were quantified. a-wave reflects (i.e., photoreceptor response) and b-wave inner retinal response) were evaluated. Bars represent mean values ± SEM. n = 4-6 for each genotype. Unpaired t-test: *P<0.05.

Three months after AAV injection, a quantitative expression analysis via qPCR was performed on purified Müller cells. A significant upregulation of Nr3c1 was noted in AAV-injected controls and a trend towards upregulation in db/db mice (**Fig. 8C**). Note that a slight Gfap upregulation as a consequence of AAV injection was still present irrespective of genotype that was not observed in the PBS sham-injected eyes, whereas no effect was observed on the expression of glutamine synthetase (Glul) (**Fig. 8C**).

Next, we asked whether this Müller cell-restricted up-regulation of GR influences the microglial response. As in the initial validation of the control versus db/db mice at 6 months of age, there was a trend of slightly higher microglial numbers in the diabetic retina of eyes that only received PBS as sham control compared to non-diabetic control eyes that were also PBS-sham injected (**Fig. 8D**). With respect to microglial soma area size, no significant difference was observed. This confirms that microglia in the retina of 6-month-old mice are just about to get activated and did not yet develop the full-blown phenotype of activated microglia yet. Of note, microglia in sham-treated diabetic mice had significantly shorter processes and, thus, occupied much smaller territories (**Fig. 8D**).

In a final step, ERG measurements were conducted to test whether GR overexpression not only alters the gene expression signature of Müller cells and microglial activation, but also affects neuronal function. Similar to in our initial characterization of untreated db/db mice (**Fig. 2**), a reduced light response (a- and b-wave) in the retinae of diabetic mice compared to controls was observed when comparing the PBS sham-injected eyes of each genotype (**Fig. 8E**). Notably, the treatment effect was most pronounced for rod-driven b-waves compared to cone-driven responses in db/db mice (**Fig. 8E**).

## Discussion

### Early functional, but late onset of morphological changes of distinct cell types in the db/db retina

The re-evaluation of the retinal phenotype of our db/db strain ensured that we had captured the right time window for our multiomics approach designed to identify early glial changes before fundamental neurodegeneration has occurred. In db/db animals, the onset of pericyte loss at 3 months of age has been described as a key feature of DR (Midena et al. 1989). We observed a decrease in the number of PDGFRβ-positive pericytes in db/db retinae, but not in animals younger than 6 months of age. Signs of microglial activation were also observed later (at 6 to 9 months of age) than published by others who found proinflammatory microglia in 2-month-old db/db mice (Arroba et al. 2016), in sum suggesting that the phenotype in our strain develops more slowly than described in the literature.

Little is known about changes of Müller cells in the course of DR progression in db/db mice. An upregulation of glial fibrillary acidic protein (GFAP) in Müller cells, as a marker of their gliotic activation, has been described in 2-month-old db/db mice (Bogdanov et al. 2014). In line with this, we demonstrate a moderate but significant upregulation of GFAP in 6-month-old animals via cell type-specific RNA-seq, qPCR, and proteomic data, whereas we were unable to detect the protein in Müller cell processes by immunolabeling. Our MACS approach has the limitation that Müller cells may be slightly contaminated with astrocytes, which express high levels of GFAP in the homeostatic retina. Thus, the increase in GFAP levels detected in the MACS-purified glial population may also be due to expression changes in astrocytes (Church et al. 2023) or may be too low to be detected in Müller cells by immunostaining. However, using physiological measurements, we were able to demonstrate that Müller cell gliosis is present in the retinae of 6- month-old db/db animals at the functional level. As we have shown previously in diabetic rats (Pannicke et al. 2006, Krugel et al. 2011), also Müller cells from db/db mice had a reduced potassium conductance and consequently a diminished ability to compensate for osmotic stress. This finding implies that the Müller cells’ capacity to perform potassium siphoning key for the retinal ion and volume homeostasis (Bringmann et al. 2006) is already perturbed in this early stage of DR and could be one driving factor of disease progression as neuronal functional and retinal tissue integrity rely on this glial housekeeping function (Reichenbach et al. 2013, Pannicke et al. 2014). Therefore, we also evaluated neuronal survival and function. As with the glial responses, which indicate a delayed onset of DR pathology in our db/db mouse strain, neurodegeneration was less pronounced at different stages compared to findings in the literature (Bogdanov et al. 2014), which describe neurodegeneration as early as 2 months of age. However, even though they seem to show a slightly reduced retinal thickness at this early age, they do not show a loss of neuronal cells over the course of aging from 2 months to 6 months, which one would expect to see in a progressive disease like DR. In our hands, only cone numbers showed significant reduction, which is in line with other reports that found predominant cone photoreceptor dysfunction in early-stage DR in zebrafish (Alvarez et al. 2010) and humans (Cho et al. 2000, McAnany et al. 2019). In agreement with other reports (Bogdanov et al. 2014, Di et al. 2019), we were able to show a functional decline of db/db retinal neurons as determined by ERG recordings in 6-month-old mice. Thus, dysfunctional Müller glia and the resulting partial disruption of retinal tissue homeostasis, which we aim to target with our glia-centric treatment approach before irreversible neuronal loss has occurred, may be an underlying cause of the observed neuronal deficits.

### Early changes in the expression landscape of Müller glia in the db/db retina

In order to identify targets for potential treatment, we determined the expression landscape of Müller cells the diabetic retina from 6-month old mice. Multiple changes as compared to cells from age-matched controls consistent with the observed functional alterations were observed in the glial mRNA and protein expression profiles. (i) The metabolism of the cells was significantly altered, with a reduction in triglyceride ligase expression, but enhanced expression of enzymes involved in mitochondrial beta-oxidation. (ii) Glial stress defence mechanisms were upregulated, such as glutathione transferase or aldehyde dehydrogenase activity that are key to aldehyde detoxification including protection from lipid peroxidation that is known to be elevated in diabetes because of enhanced levels of oxidative stress (Pan et al. 2008, McDowell et al. 2016, Augustine et al. 2020). (iii) Signaling pathways, including those mediated by TGFβ, insulin-like growth factor-both via receptor tyrosine kinase-related mechanisms-and in addition G protein-mediated signaling in general, were upregulated in Müller cells from diabetic mice. The latter seems in agreement with a recent finding that modulation of G-protein-mediated signaling by application of phosphodiesterase inhibitors is beneficial in various models of retinal degeneration and enhances the tissue stress resilience (J et al. 2023). (iv) Several potassium channels and transporters were downregulated especially at protein level. At the early stage of DR that we studied here, the overall response of Müller cells is well suited to preserve and protect the retinal tissue, although initial functional adaptations (e.g., reduced potassium conductance, impaired cell volume regulation, upregulation of GFAP) indicate that the cells potentially are on the verge of developing a less supportive, gliotic phenotype.

Previous bulk RNA-seq or proteomic approaches performed on 10-12 week-old db/db or Akimba (Ins2Akita×Vegfa+/-) mice, a type 1 diabetes model of DR, identified GO terms related to synaptic transmission, glutamate transport and mitochondrial genes as downregulated. Pathways related to a metabolic shift from glycolysis to oxidative phosphorylation, activation of microglia/macrophages, metal ion and oxidative stress response were upregulated transcripts (Bogdanov et al. 2014, Van Hove et al. 2020) or proteins (Ly et al. 2014), largely confirming our data. Kandpal et al. (Kandpal et al. 2012) investigated mRNA changes by whole retinal RNA-seq 8 months after diabetes induction in a streptozotocin model of type I diabetes in mice and found alteration in pathways such as inflammation, microvasculature remodelling, apoptosis, glucose metabolism, Wnt signaling and photoreceptor biology indicating a more advanced stage of disease. In addition, Grant et al. (Grant et al. 2004) showed that ischemia-mediated overexpression of growth factors such as vascular endothelial growth factor, insulin-like growth factor-1, angiopoetin-1 and -2, stromal-derived factor-1, fibroblast growth factor-2, and tumor necrosis factor occurs in DR. This is consistent with our finding that growth factor signaling pathways are upregulated in Müller cells.

However, when comparing our data with those of studies performed on whole retinal extracts, the observed discrepancies regarding differential regulation of genes involved in metabolism, homeostasis and inflammation could be due to a response of different cell types. Single cell RNA-seq (scRNA-seq) of retinal tissue from 3-month-old Akimba mice identified upregulation of genes associated with ribosome, cytoskeleton, immune system processes, S100 proteins, glutathione metabolism, iron ion homeostasis, cell cycle regulation/apoptosis, and oxidative phosphorylation (OXPHOS) networks in Müller cells/macroglia (Van Hove et al. 2020). These findings are in agreement with our data from 6-month-old db/db mice. For example, genes involved in glutathione metabolism such as Gstm1 were upregulated in macroglia from Akimba mice as well as specifically in Müller cells in our diabetes model, indicating a consistent anti-stress response of the cells in both models. The same accounts for Aldh1l1, which was identified in the van Hove data set and was also clearly up-regulated at transcript and protein level in our bulk data on Müller cells. Moreover, van Hove et al. (2020) reported that genes involved in glycolysis, central nervous system development, and OXPHOS were downregulated in Akimba macroglia. Some of these genes were also significantly expressed by Müller cells, as well as down-regulated in the diabetic retina in our data set: GUF1 homolog, GTPase (Guf1) is involved in metabolic processes of organo-nitrogen compounds (**Tab. S1**). Fibroblast growth factor binding protein 3 (Fgfbp3) and phosphotyrosine interaction domain containing 1 (Pid1) contribute to the regulation of phosphate metabolism (**Tab. S1**).

Importantly, our study differs from previously published work in that we provide cell type-specific insights into disease-associated expression profiles not only at the transcript level but also, for the first time, at the protein level. The importance of this for delineating the functional change of a cell is underscored by the fact that we, like many others before us (Vogel et al. 2012, Liu et al. 2016, Kjell et al. 2020), found considerable discordance in the transcript and protein levels of distinct genes (e.g. YAP1) - so one cannot reliably interpolate from changes in transcript levels to how the corresponding protein expression will change. Since in most cases it is the protein that mediates the actual function of a (protein-coding) gene, we consider this multi-layered omics approach is much more informative for understanding how diabetes affects cellular interactions in the retina than studies that focus solely on mRNA analysis.

### The glucocorticoid receptor as potential master regulator of Müller cells in the diabetic retina

In our search for key regulators that drive Müller cell changes in DR progression that could then be targeted by a therapeutic approach, the glucocorticoid receptor (GR, gene ID: Nr3c1) was identified as one of the most promising candidates for several reasons. It was the one of few transcription factors for which ATAC-seq performed on purified Müller cells identified a significant alteration in the accessibility of its DNA-binding motives. In line with that, also the oPOSSUM-3 transcription factor binding site cluster analysis on the basis of the RNA-seq data of differentially expressed genes in Müller cells from db/db mice, besides others, identified the GR as one of the most likely transcription factors that could be causative for the observed changes in gene expression profiles. Finally, only the GR fulfilled our additional screening criteria – a high expression in Müller cells and a differential expression in Müller cells of 6-month-old db/db mice at transcript and protein level. Importantly, we were able to clearly localize GR immunoreactivity to Müller cell nuclei, suggesting that it is active in the cells, which is consistent with the high corticosterone levels present especially in the blood plasma of db/db animals together with the increased level of GR phosphorylation determined by Western blot analysis. Indeed, increased GR mRNA expression within 4 h after NMDA-induced damage, normalized to pre-damage levels by day 2 and below pre-damage levels by day 3, has also been described in the chicken retina (Gallina et al. 2014). A similar response, but with much longer time scales, may also be active during DR progression.

Even though recent studies suggest potential roles for the GR in DR, it is important to note that the understanding of exact mechanisms of the interplay between the GR, inflammation, vascular changes, neuroprotection, and other molecular pathways involved in the development and progression of DR are still being elucidated. DR is associated with chronic low-grade inflammation and immune dysregulation, which could also be confirmed in our present study. The GR has been implicated in regulating vascular permeability in the retina, e.g by inhibiting VEGF-induced permeability in endothelial cells, potentially offering a protective effect against retinal vascular leakage (Imai et al. 2017). In human retinal endothelial cells (HRECs), dexamethasone, a GR ligand, significantly repaired glucose-induced cell loss and enhanced permeability, suggesting a potential role of GR in protecting this cell type in high glucose condition and facilitating endothelial cell repair in the diabetic retina (Stewart et al. 2016, Sulaiman et al. 2018). The GR has been associated with neuroprotective effects in various models of neurodegeneration. Activation of the GR has shown to promote neuronal survival and attenuate apoptotic cell death, suggesting a potential role in protecting retinal neurons from DR-induced damage (Wenzel et al. 2001, Wenzel et al. 2003, Zhang et al. 2013).

The major ligand of the GR in mice is the glucocorticoid corticosterone. Circulating levels of glucocorticoids are regulated by the hypothalamic-pituitary-adrenal (HPA) axis. During stress, the HPA axis is activated and an increase in glucocorticoids helps the body cope with and recover from the stressful situation (Gjerstad et al. 2018). As in our study, others have described elevated corticosterone levels in mouse models of type 1 and type 2 diabetes (Erickson et al. 2017). Similarly, patients with diabetes present with higher urinary free cortisol (Roy et al. 1998). Chronic administration of glucocorticoids can constitutively downregulate GR expression via an autoregulatory loop (Vandevyver et al. 2014). Therefore, persistently high concentrations of corticosterone in the blood may negatively regulate GR expression in diabetic mice.

As a nuclear hormone receptor, GR coordinates inflammation, cell proliferation and differentiation in target tissues. Consistent with our findings, Gallina et al. (2014) found that GR was mainly located in Sox2-positive nuclei of Müller cells in mouse, guinea pig, dog, and human retinae. Partial loss of retinal GR results in a thinner INL and that it plays a critical role in maintaining retinal homeostasis by regulating the inflammatory response (Kadmiel et al. 2015). Furthermore, Gallina et al., 2015 found that activation of GR inhibited the reactivity of microglia and the loss of retinal neurons upon excitotoxic tissue damage. The activated GR can regulate the expression of target genes through multiple ways. It can lead to transactivation or transrepression of gene transcription directly by binding to glucocorticoid-response elements (GREs) in regulatory regions of specific target genes (Sulaiman et al. 2018, Ghaseminejad et al. 2020). Furthermore, the GR can act through protein-protein interactions with other TFs like the nuclear factor kappa-light-chain enhancer of activated B cells (NF-κB), activating protein-1 (AP-1) or YAP1 (Wenzel et al. 2003, Sulaiman et al. 2018, Ghaseminejad et al. 2020).

YAP1, for instance, is the central factor on which the Hippo pathway converges and that coordinates cell proliferation and stemness. In case the Hippo signaling pathway is less active, YAP1 translocates into the nucleus, followed by the activation of its downstream targets which are associated amongst others with increased proliferation and cell cycle entry (Totaro et al. 2018, Ma et al. 2019). In line with this, inhibition of YAP1 signaling via the Hippo pathway prevents Müller cell proliferation upon injury in mammals whereas proliferation is induced via enhanced YAP1 activation (Hamon et al. 2019, Rueda et al. 2019). Interestingly, YAP1 seems to be co-regulated with GR (Sorrentino et al. 2017, Kim et al. 2020), which we could confirm at transcript level. However, at protein level, we detected higher YAP1. One reason could be that the high levels of glucose lead to production of uridin-5′-diphospho-N-acetylglucosamin (UDP-GlcNAc) and this results in increased YAP O-GlcNAcylation which can stabilize YAP protein levels (Peng et al. 2017, Zhang et al. 2017). Therefore, the ultimate impact of changing GR levels on YAP1-mediated pathways should be investigated in more detail in future studies.

The GR target gene cluster identified by the oPOSSUM-3 analysis consisted mainly of downregulated genes. Interestingly, a quarter of the significantly downregulated genes in Müller cells of the db/db mouse retina were also potential target genes of the GR. One such example is the transcript of forkhead box o1 (Foxo1). It is involved in the control of insulin sensitivity, hepatic glucose production, and blood glucose levels (Wu et al. 2018). Carbonic anhydrase 4 (Ca4), a member of a large family of zinc metalloenzymes that catalyzes the reversible hydration of carbon dioxide, is another example of a significantly downregulated GR target gene in Müller cells of diabetic mice. Ca4 is essential for acid removal from the retina, while mutations in Ca4 impair pH regulation and cause retinal photoreceptor degeneration (Yang et al. 2005). The strong impact of these two exemplary GR target genes on retinal integrity suggests that regulation of the GR target gene cluster in the diabetic retina has great potential to slow disease progression.

### AAV-vectored GR overexpression in Müller cells improves neuronal function in db/db mice

Treatment strategies for DR (e.g. anti-VEGF, intravitreal steroids, photocoagulation) require lifelong repeated invasive interventions with significant side effects. Given our findings that the GR is specifically regulated in Müller cells of the diabetic retina and that its activity has beneficial effects on the cells as well as on neighbouring neurons, we tested our hypothesis that the GR is a promising potential target for a novel gene therapeutic approach. We restricted GR overexpression to Müller cells to ensure its specific action in glial cells alone, leaving unaltered retinal neurons, which may be less able to cope with exogenous gene overexpression. This also ensured that other ocular tissues, which typically experience unwarranted side effects such as the formation of lens cataracts with the continued use of intravitreal steroids, were spared (Wong et al. 2016). After validating the successful overexpression of GR in Müller cells, we analyzed the Müller glial response, but found no difference, e.g. in the upregulation of the gliosis marker GFAP, which was still slightly upregulated in AAV-injected eyes of both genotypes 3 months after injection. This persistent moderate gliotic response of the cells after transduction by the virus was not observed in sham-injected eyes receiving PBS only. This finding underscores the current debate in the field that a thorough re-evaluation of the immunogenic potential of AAV serotypes and expression constructs used is necessary to optimize the efficacy of gene therapy approaches (Verdera et al. 2020, Arjomandnejad et al. 2023).

We found a moderate but significant effect on microglia upon overexpression of GR in Müller cells. As in the initial validation of the retinal phenotype in the 6-month-old db/db mice, we did not observe genotype-dependent differences in microglia number or soma area. However, we additionally analyzed the area occupied by microglial processes, which shrinks upon microglial activation (Scholz et al. 2015, Mages et al. 2019). Indeed, we observed a significant reduction of the microglial occupied area in the diabetic retina compared to controls, which was reverse by overexpression of GR in Müller cells. This is a first indication supporting the idea that GR activity in Müller cells may also influence their close cross-talk with microglia, which is central to modulate retinal immune homeostasis (Wang et al. 2014, Kaczmarek-Hajek et al. 2018, Mages et al. 2019, Pauly et al. 2019, Schmalen et al. 2021).

Finally, we used ERG recordings as a highly sensitive readout to test for potential treatment effects on neuronal function, as we found significant alterations db/db mice at 6 months of age, whereas we found very little morphological changes at this stage of the disease, so that treatment effects would have been difficult to assess by morphometry. We confirmed the significantly reduced retinal responsiveness to light in the diabetic retina also when comparing the PBS sham-treated eyes of both genotypes. Overexpression of GR in Müller cells resulted in a moderate improvement, especially in the cells integrating the input from rod photoreceptors, as reflected by the most pronounced differences between b-waves measured under scotopic conditions. This suggests that inner retinal cells, such as bipolar, amacrine or ganglion cells, may benefit most from the changes associated with GR overexpression in Müller cells, which would be consistent with our recent study showing that improved homeostatic functions of Müller cells, such as a better maintained potassium conductance, are specifically relevant to this cell population (Pannicke et al. 2014, Mages et al. 2019). Our findings demonstrating that targeting Müller cells alone is sufficient to achieve beneficial treatment effects in the diabetic retina are in line with the recent study by Niu et al. (Niu et al. 2021). They used AAV to overexpress RLBP1, a Müller cell protein that acts as a soluble retinoid carrier essential for proper photoreceptor function in the visual cycle, and also achieved significant treatment effects.

## Conclusion

We identified the GR as a promising candidate to be targeted by gene therapeutic approaches as (i) we and others demonstrated GR expression to be almost exclusive to Müller glia, in a highly conserved manner amongst warm-blooded vertebrates (i.e. retinae of chicks, mice, guinea pigs, dogs and humans), (ii) many transcription factors downstream or co-active with GR are specifically expressed in Müller glia in the retina, (iii) GR expression is significantly and specifically down-regulated in gliotic Müller cells of the diabetic mouse retina, (IV) the transcription factor binding site cluster analysis (oPOSSUM) based on our RNA sequencing data identified GR as one of few strong candidates to explain the diabetes-associated expression changes in Müller cells which was also confirmed by Atac-seq, (v) stimulating GR signaling using cortisol in retinal explant cultures fostered the expression of genes important for neuron-supportive Müller cell functions, (vi) transgenic mice receiving injections of the selective GR agonist triamcinolone acetonide (TA) 4 days prior to conditional Müller cell ablation presented with significantly higher Müller cell survival and if left untreated, loss of Müller cells caused photoreceptor degeneration, vascular leakage and intraretinal neovascularization – hallmarks of DR, (vii) previous studies have shown that TA reduces vascular leakage, inhibits the secretion of VEGF and prevents osmotic swelling of Müller cells and (viii) finally, treatment with GR agonists (e.g. TA) has proven to be effective in inflammatory diseases, DR included, as it counterbalances typical changes associated with DR such as breakdown of the blood retinal barrier or onset of neuroinflammation. Future studies are needed to optimize AAV-vectored GR delivery and to tune its activity levels in Müller glia to avoid potential adverse cellular responses to AAV-driven construct expression, with the overall goal of improving the therapeutic efficacy of the approach presented here with initial pilot data.

## Methods

### Animals

Db/wt heterozygous mice (BKS.Cg-Dock7m^+/+^Leprdb/J) were obtained from Jackson Laboratories (https://www.jax.org/strain/000642) and BKS-Lepr^db/db^/JOrlRj were obtained from Janvier Labs (https://www.janvier-labs.com/en/fiche_produit/diabetique_mouse/) and maintained in our animal facility. All experiments were performed in accordance with European Community Council Directive 86/609/EEC and were approved by local authorities (ROB-55.2-2532.Vet_02_18_20). Animals had free access to water and food in a climate-controlled room with a 12-h light-dark cycle. Mice of both sexes at 12, 24, and 38 weeks of age were used for the experiments. The genotyping of the mice was performed based on the protocol previously published by (Peng et al. 2018) using the following primers: fw_i_ -ATT AGA AGA TGT TTA CAT TTT GAT GGA AGG; fw_o_ - TTG TTC CCT TGT TCT TAT ACC TAT TCT GA; rev_i_ - GTC ATT CAA ACC ATA GTT TAG GTT TGT CTA; rev_o_ - CTG TAA CAA AAT AGG TTC TGA CAG CAA C. PCR of DNA from wildtype mice revealed two bands at 610 and 264 bp, that from db/+ mice three bands at 610, 406, and 264 bp, and that from homozygous db/db mice two bands at 610 and 406 bp. Wild type and db/+ mice were used as controls as they do not develop the diabetes phenotype.

### Corticosterone level measurement in mouse blood

A corticosterone ELISA kit (Abcam, Cambridge, UK) was used according to the manufacturer’s recommendations to measure corticosterone levels in the blood plasma of five 6-month-old mice per genotype. Samples were collected in blood collection tubes, centrifuged at 3000g for 10 minutes at 4°C, and stored at -20°C. Blood plasma samples were used for ELISA at a dilution of 1:100.

### Retinal explant culture

Retinal explants from wild-type mice were cultured for two or five days. Mice were euthanatized, eyes enucleated, and retinae carefully removed. Explant cultures were placed in a 24-well plate containing 500 μL medium (DMEM/F-12, GlutaMAX™, 1:100 antibiotic-antimycotic; Thermofischer) per well. Retinae were placed in the center of a Whatman® Nuclepore™ track-etched membrane (Merck, Darmstadt, Germany) and covered with a drop of medium. The retinal explants were maintained at 37°C in 5% CO2 with daily medium changes, while cortisol was added twice daily to maintain a constant high concentration of 500 ng/ml.

### Patch-clamp recordings of single Müller cells

For whole-cell patch-clamp experiments, cells were isolated as described above. The cells were stored at 4 °C in serum-free minimum essential medium until use within 4 h after cell isolation. Müller cells were identified according to their characteristic morphology. The currents were recorded at room temperature using the Axopatch 200A amplifier (Axon Instruments, Foster City, CA, USA) and the ISO-2 software (MFK, Niedernhausen, Germany). The signals were low-pass filtered at 1 or 6 kHz (eight-pole Bessel filter) and digitized at 5 or 30 kHz, respectively, using a 12-bit A/D converter. Patch pipettes were pulled from borosilicate glass (Science Products, Hofheim, Germany) and had resistances between 4 and 6 MΩ when filled with a solution containing (mM): 10 NaCl, 130 KCl, 1 CaCl2, 2 MgCl2, 10 EGTA, and 10 HEPES, adjusted to pH 7.1 with Tris. The recording chamber was continuously perfused with ECS. To evoke membrane currents, de- and hyperpolarizing voltage steps of 250 ms duration, with increments of 10 mV, were applied from a holding potential of -80 mV. The amplitude of the steady-state inward currents was measured at the end of the 250-ms voltage step from -80 to -140 mV. The membrane capacitance of the cells was measured by the integral of the uncompensated capacitive artifact (filtered at 6 kHz) evoked by a 10-mV voltage step in the presence of extracellular BaCl2 (1 mM). Current densities were calculated by dividing inward current amplitudes evoked by 60 mV hyperpolarization by the membrane capacitance. The resting membrane potential was measured in the current-clamp mode. Data are expressed as mean ± standard deviation (SD), significance was determined by the non-parametric Mann-Whitney U test.

### Measurement of Müller cell volume regulation

Volume changes in retinal Müller cells were measured as described (Slezak et al. 2012). Briefly, retinal slices were loaded with the vital dye Mitotracker Orange (10 µM, excitation: 543 nm, emission: 560 nm long-pass filter; Life Technologies), which is preferentially taken up by Müller cells (Uckermann et al. 2004). Slices were exposed to hypotonic solution (60% of control osmolarity using distilled water) for 4 min with or without test substances. Somata of labelled Müller cells were imaged using confocal microscopy (custom-made VisiScope CSU-X1 confocal system equipped with high-resolution sCMOS camera; Visitron Systems, Puchheim, Germany) and their cross-sectional areas were measured (ImageJ).

### Histological and immunohistochemical staining

Enucleated eyes were immersion-fixed (4% paraformaldehyde (PFA) for 1 h, room temperature), washed with phosphate-buffered saline (PBS), cryoprotected in sucrose, embedded in Cryomatrix^TM^ (Thermofisher, Germany) and cut in 10 µm sections using a cryostat. Similarly, retinal explant cultures were removed from the medium and were fixed still directly on culture inset with 4% PFA at for 1 h at room temperature. Retinal sections were permeabilized (0.3% Triton X- 100 plus 1.0% DMSO in PBS) and blocked (3% DMSO, 0,1 % Triton X-100, 5 % goat oder donkey serum in PBS) for 1 h at room temperature. Primary antibodies (**Tab. 1**) were incubated overnight at 4°C. Sections were washed (1% bovine serum albumin in PBS) and incubated with secondary antibodies (2 h at room temperature; **Tab. 1**). Cell nuclei were labeled with DAPI (1:1000; Life Technologies). Sections were mounted using Aqua-Poly/Mount (Polysciences Europe, Eppelheim, Germany). Retinal flatmounts were labeled using a similar protocol, except that tissue was permeabilized by higher concentrations of 2% BSA, 0.5% triton und 5% goat/donkey serum in PBS and secondary antibodies were also incubated at 4°C overnight. Control experiments without primary antibodies showed no unspecific labeling. Images were taken with custom-made VisiScope CSU-X1 confocal system (Visitron Systems, Puchheim, Germany) equipped with high-resolution sCMOS camera (PCO AG, Kehlheim, Germany). For cell number quantification, images of retinal sections or z-stacks of retinal flatmounts were taken scanning through the nerve fiber layer to the outer nuclear layer of flatmounts (optical section thickness: 1 µm). Scans were confined to the central retina in close proximity to the optic nerve head, and morphometric parameters were assessed using Fiji Schindelin (Schindelin et al. 2012).

**Table 1.**
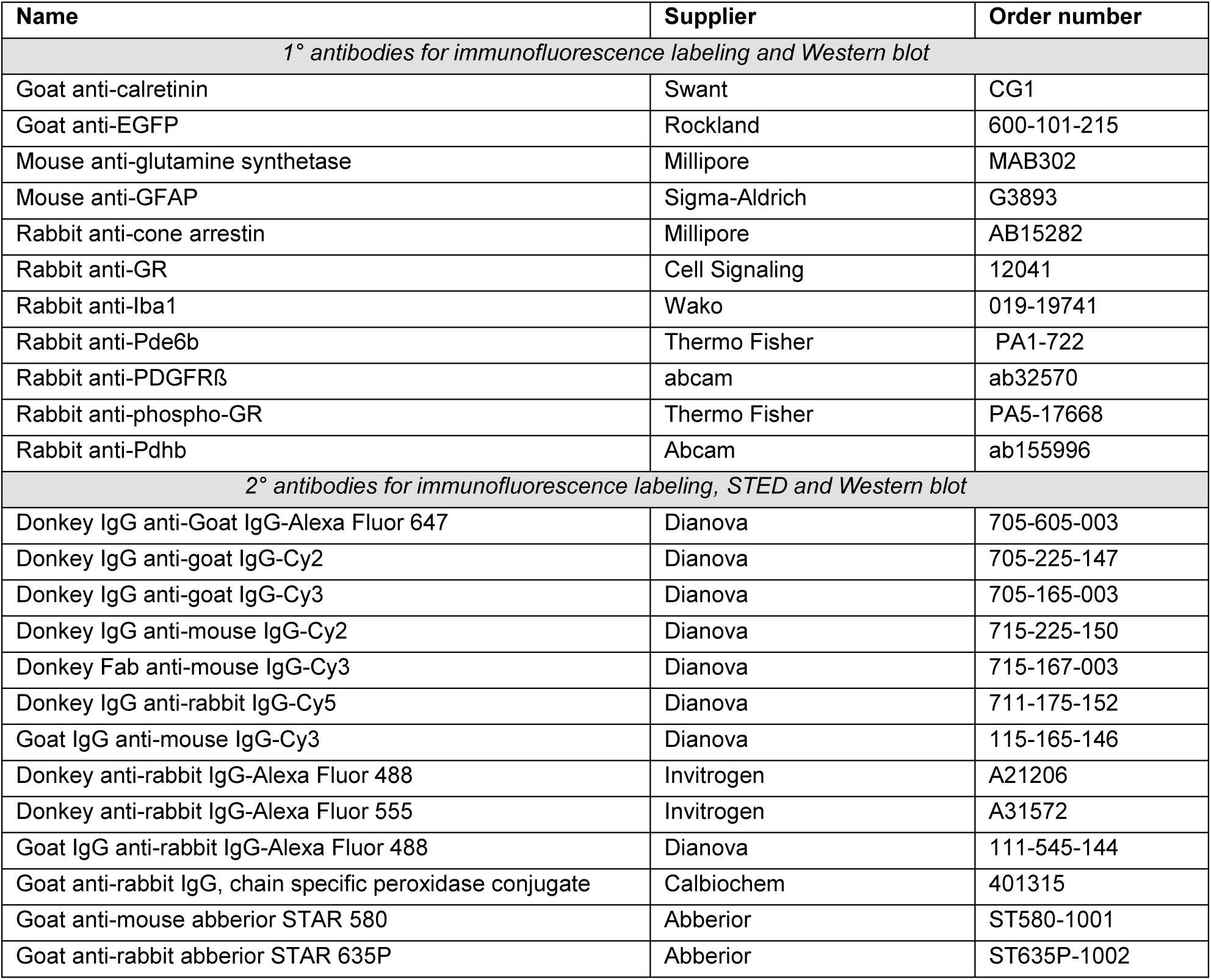
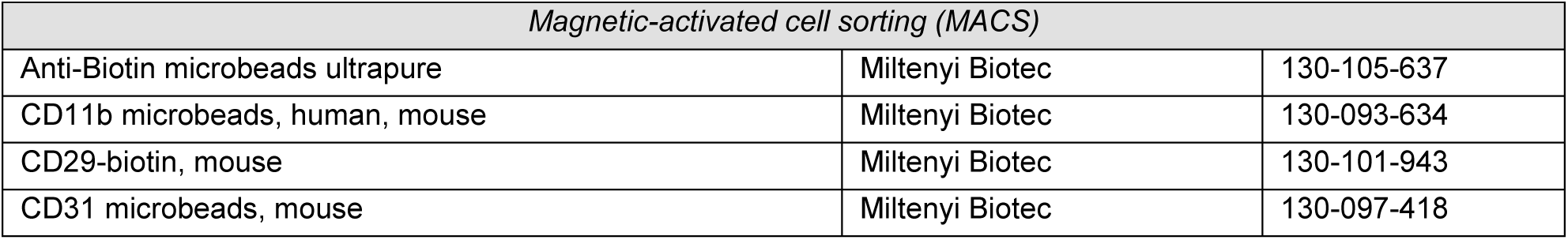
Antibodies and antibody-conjugated beads used in the present study.

For super resolution microscopy, fixed retinal sections were permeabilized with Triton X- 100 (0.5 % in 2% BSA in PBS) for 2h and incubated with primary antibodies in blocking solution overnight at 4°C. After rinsing several times with PBS, sections were incubated with secondary antibodies (Abberior Star) and DAPI for 2h in blocking solution at RT, subsequently rinsed 3 times with PBS and once with distilled water. Sections were mounted with ProLong Gold antifade reagent (Invitrogen, Life Technologies). Stimulated emission depletion (STED) microscopy was performed at the Core Facility Bioimaging of the Biomedical Center of the LMU München with an inverted Leica SP8 STED X WLL microscope using appropriate lasers for fluorophore excitation (405 nm; pulsed white light laser 470 – 670 nm). Image acquisition was performed with a 93x/1.3 NA glycerol immersion objective with the pixel size set to 32 nm. To avoid bleed-through, the color channels were recorded sequentially. The following spectral settings were used: DAPI (excitation: 405 nm; emission: 415 – 471 nm; PMT), AbberiorStar 580 (580 nm; 590 – 620 nm; HyD; depletion laser: pulsed 775 nm, at 20% intensity), AbberiorStar 635P (635 nm; 645 – 702 nm; HyD; depletion laser: pulsed 775 nm, at 12% intensity). Image contrast and brightness was adjusted with the open source software FIJI (Schindelin et al., 2012).

### Trypsin digest

The vascular architecture of the retina was studied using a trypsin digest-based protocol as described recently (Chou et al. 2013). Enucleated eyes were fixed in 4% PFA for 24 hours at 4°C before retinal isolation. The cornea and lens were removed, and the retina was carefully separated from the sclera and choroid. After five gentle shaking washes with sterile water at RT for 30 minutes, the retinae were left in water on the shaker at RT overnight. The water was removed and the retinae were digested with 3% trypsin in 0.1 M Tris buffer with gentle shaking at 37°C for 1.5h. The trypsin was replaced with sterile water and the internal limiting membrane was removed. The retinal vasculature was then separated from the rest of the tissue by a series of 5-minute water washes. When little or no debris remained, the vascular network was carefully transferred to a microscope slide using a trypsin-coated, fire-polished glass pipette. After drying on a hot plate at 37°C overnight, hematoxylin and eosin (H&E) staining was performed. The specimens were then mounted using Aqua-Poly/Mount (Polysciences Europe) and a custom-built VisiScope CSU-X1 confocal system (Visitron Systems) equipped with a bright field unit. The number of endothelial cells, pericytes, and acellular capillaries was assessed using the cellcounter tool of Fiji (Schindelin et al. 2012), and endothelial cells and pericytes were identified by their characteristic morphology (e.g., their nuclei) and their location on the vessel wall. The calculation of the endothelial cell/pericyte ratio is described by Midena et al. (Midena et al. 1989).

### Magnetic activated cell sorting

Different cell types of the retina were sequentially separated by magnetic activated cell sorting (MACS) as previously described (Pauly et al. 2019). Retinae were isolated and digested with 0.2 mg/ml papain in PBS/Glucose at 37 °C for 30 min. Afterwards, three washing steps with PBS/Glucose were performed and the tissue incubated with 200 U/ml DNase I at RT for 4 min. The retinae were dissociated in extracellular solution (ECS) to get a single retinal cell solution and centrifuged at 600 g and 4 °C for 10 min. The supernatant was removed and the cells were resuspended in ECS and incubated with CD11b microbeads (**Tab. 1**) which have been developed for the positive isolation of primary mouse CD11b^+^ microglia at 4 °C for 15 min. After centrifugation at 600 g and 4 °C for 10 min the cells were resuspended in ECS and transferred onto a large cell column using a fire polished glass pipette. The cells were separated according to the manufacturer’s recommendation. The CD11b^+^ fraction was centrifuged and the cells stored at -80°C. The CD11b^-^ cells (flow through) were centrifuged, resuspended in ECS and incubated with CD31 (Pecam1)-microbeads (**Tab. 1**) for the positive selection of CD31^+^ endothelial cells at 4 °C for 15 min. After centrifugation at 4 °C and 600 g for 10 min the cells were resuspended in ECS, transferred onto large cell columns and eluted according to the manufacturer’s recommendation. The CD31^+^ fraction was centrifuged and the cells stored at -80 °C. The CD31^-^ cells (flow through) were centrifuged, resuspended in ECS and incubated with CD29-biotin at 4 °C for 15 min. The antibody binds to CD29 (**Tab. 1**), also known as integrin ß1, which is a cell-surface receptor expressed by Müller cells. Retinal astrocytes are assumed to be found in a small amount in the CD29^+^ fraction. The cells were centrifuged at 4 °C and 600 g for 10 min, resolved in ECS and incubated with Anti-Biotin microbeads which enables the positive selection of the CD29^+^ Müller cells at 4 °C for 15 min. After centrifugation the cells were resuspended in ECS, transferred onto the columns and separated according to the manufacturer’s recommendation. The remaining CD29^-^ fraction mostly contained the retinal neurons. The remaining samples were centrifuged at 10000 g and 4 °C for 15 min and the cell pellet was stored at -80 °C.

### qRT-PCR

Total RNA was isolated from enriched cell populations using the PureLink^®^ RNA Micro Scale Kit (Thermo Fisher Scientific, Schwerte, Germany). A DNase digestion step was included to remove genomic DNA (Roche, Mannheim, Germany). First-strand cDNAs from 10-50 ng of total RNA were synthesized using the RevertAid H Minus First-Strand cDNA Synthesis Kit (Fermentas by Thermo Fisher Scientific, Schwerte, Germany). Primers were designed using the Universal ProbeLibrary Assay Design Center (Roche, **Tab. 2**). Transcript levels of candidate genes were measured by qRT-PCR using cDNA with the QuantStudio 6 Flex Real-Time PCR System (Life Technologies, Darmstadt, Germany) according to the company’s guidelines.

**Table 2.**
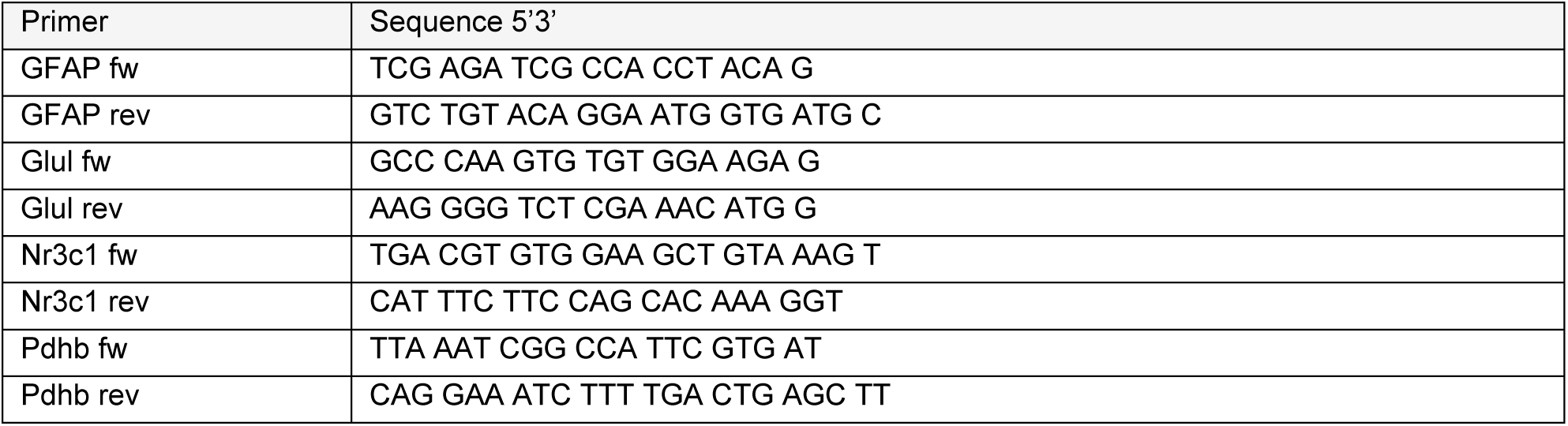
Primers used for qPT-PCR.

### Western blot

Gels containing 12% acrylamide were used for sodium dodecyl sulfate-polyacrylamide gel electrophoresis (SDS-PAGE). Samples were mixed with 5X SDS-PAGE sample buffer, incubated at 95°C for 10 minutes, and loaded onto the gel next to the prestained Protein™ ladder plus (Thermofisher Scientific, Darmstadt, Germany). Electrophoresis was performed at 60 V for 45 min and at 150 V for 70 min. The proteins were then transferred to a 0.22 µm PVDF membrane by blotting at 24 V for 35 min using a Trans-Blot Turbo^TM^ system (BioRad, Feldkirchen, Germany). The membrane containing the proteins was blocked in 5% BSA in TBST at RT for 1 h to avoid unspecific binding of the antibodies. The expected size of GR was 95 kDa and that of phosphorylated GR was 86 kDa. PDHB was used as a housekeeping protein because it is present at similar levels in all retinal cell types. Since the expected size was 35 kDa and the molecular weight of the phospho-/GR was 86/95 kDa, the membranes were cut into two pieces at the 70 kDa band of the standard. The membrane pieces were shaken in the primary antibody overnight at 4°C. The next day, they were washed three times in TBST for 10 minutes and incubated with the secondary antibody in 5% BSA in TBST at RT for 2 hours. The Clarity Max™ Western ECL Substrate Kit (BioRad) was used and the bands were visualized using the ChemiDoc XRS+ system (BioRad). GelAnalyzer 19.1 (www.gelanalyzer.com) was used to quantify the bands.

### Bulk-RNA sequencing

After immunoseparation of retinal cell types, total RNA was isolated from cell pellets using the PureLink® RNA Micro Scale Kit (Thermo Fisher Scientific, Schwerte, Germany). Validation of RNA integrity and quantification was performed using the Agilent RNA 6000 Pico chip analysis according to the manufactures instructions (Agilent Technologies, Waldbronn, Germany). Enrichment of mRNA and library preparation (Nextera XT, Clontech), library quantification (KAPA Library Quantification Kit Illumina, Kapa Biosystems, Inc., Woburn, MA, USA) as well as sequencing on an Illumina platform (NextSeq 500 High Output Kit v2; 150 cycles) were performed at the service facility KFB Center of Excellence for Fluorescent Bioanalytics (Regensburg, Germany; www.kfb-regensburg.de). After de-multiplexing, a total of at least 20 million reads per sample was reached. Quality control of the reads and quantification of transcript abundance was performed with the Tuxedo suit, as described elsewhere (Brandl et al. 2015, Grassmann 2019). Briefly, adapter sequences were removed with cutadapt (Kechin et al. 2017) and several quality control measures were queried with fastqc. No major problems with the sequencing data was detected. Next, the trimmed reads were aligned to the reference genome/transcriptome (mm10) with HISAT2 (Kim et al. 2015) and transcript abundance was estimated with stringtie, expressed as fragments per 1,000 base pairs of transcript per million reads (FPKM).

### ATAC-seq and data analysis

Müller cells were purified using the MACS protocol and subjected to the omni-ATAC protocol with slight modification (Corces et al. 2017). Briefly, 100.000 viable, pelleted cells were resupended in cold ATAC resuspension buffer (10mM Tris-HCl pH 7.5 (Invitrogen, catalog nr 15567027), 10mM NaCl (Invitrogen, catalog nr AM9760G), 5mM MgCl2 (Invitrogen, catalog nr AM9530G), 0.1% IgePAL (Sigma Aldrich, catalog nr I3021-50ML), 0.1% Tween-20 (Sigma Aldrich, catalog nr P9416-50ML), 0.01% Digitonin (Promega, catalog nr G9441)) and incubated on ice for 3min. Lysed cells were washed with cold ATAC wash buffer (10mM Tris-HCl pH 7.5, 10mM NaCl, 5mM MgCl2, 0.1% Tween-20) and nuclei pelleted 10min at 500g. Nuclei were resuspended in 25uL of transposition mix (12.5ul 2x TD buffer, 1.25ul transposase (Illumina, catalog nr 20034197), 8.25ul PBS, 0.25ul 1% Digitonin, 0.25ul 10% Tween-20, 2.5ul nuclease-free water (Invitrogen, catalog nr AM9937)) and incubated at 37°C for 30min, 1000RPM. Transposed fragments were purified using a Zymo Clean and Concentrator5 kit (Zymo, catalog nr R1013), followed by amplification (Ultra II Q5, NEB, catalog nr M0544L) and double size selection of 100-1000bp fragments with AMPure XP beads (Beckman Coulter, catalog nr A63881). Library was sequenced using 2×50bp reads.

Differential accessible (DA) peaks were analyzed using the csaw package (Lun et al. 2016), with loess normalization. Within DA peaks, enriched motifs were identified using the Monalisa package (Machlab et al. 2022).

### Label-free liquid chromatography mass spectrometry

MACS enriched retinal cell types from four control and diabetic mice at 24 weeks of age were collected as detailed in (Pauly et al. 2019) and proteolyzed with Lys-C and trypsin by a modified FASP procedure as described (Frik et al. 2018) resulting in two fractions after sequential proteolysis with first Lys-C (Wako Chemicals) followed by trypsin (Promega). Peptides were analyzed on a Q Exactive HF mass spectrometer (Thermo Fisher Scientific) online coupled to a UItimate 3000 RSLC nano-HPLC (Dionex). Samples were automatically injected and loaded onto the C18 trap cartridge and after 5 min eluted and separated on the C18 analytical column (Acquity UPLC M-Class HSS T3 Column, 1.8 μm, 75 μm × 250 mm; Waters) by a 90 min non-linear acetonitrile gradient at a flow rate of 250 nl/min. MS spectra were recorded at a resolution of 60000 with an AGC target of 3 × 1e6 and a maximum injection time of 30 ms from 300 to 1500 m/z. From the MS scan, the 10 most abundant peptide ions were selected for fragmentation via HCD with a normalized collision energy of 27, an isolation window of 1.6 m/z, and a dynamic exclusion of 30s. MS/MS spectra were recorded at a resolution of 15000 with a AGC target of 1e5 and a maximum injection time of 50 ms. Unassigned charges, and charges of +1 and >8 were excluded from precursor selection.

Acquired raw data were loaded to the Progenesis QI software for MS1 intensity based label-free quantification (Nonlinear Dynamics, Waters), separately for the two different proteolyzed fractions and for each cell type. After alignment in order to reach a maximum overlay of peptide features, filtering of singly charged features and features with charges > 7 and normalization to correct for systematic experimental variation, all MSMS spectra were exported and searched against the Swissprot mouse database (16772 sequences, Release 2016_02) using the Mascot search engine. Search settings were: enzyme trypsin or Lys-C, respectively, 10 ppm peptide mass tolerance and 0.02 Da fragment mass tolerance, one missed cleavage allowed, carbamidomethylation was set as fixed modification, methionine oxidation and asparagine or glutamine deamidation were set as variable modifications. A Mascot-integrated decoy database search calculated an average false discovery of < 1% when searches were performed with a mascot percolator score cutoff of 13 and significance threshold of 0.05. Peptide assignments were re-imported into the Progenesis QI software. The two datasets per cell type were combined and the abundances of the top three intense unique peptides allocated to each protein were summed up. Missing values were imputed by low abundance imputation in Perseus. The resulting normalized abundances of the single proteins were then used for calculation of fold changes of proteins and significance values by a Student’s t-test (based on log transformed values).

A principal component analysis of the proteomics data was performed and a heatmap which was focusing on proteins differentially regulated in Müller cells of diabetic mice generated by filtering p<0.05 and at least twofold difference was performed via the R Core Team 2020. The effect of cortisol treatment on retinal explants was investigated in six retinae per treatment group. Samples were processed as described before. A principal component analysis of the proteomes was performed and a heatmap generated by filtering P< 0.05 and at least 1,3fold difference. Pathway enrichment analysis for molecular functions for both data sets was done via the open-access platform PANTHER (released 20221013).

### Comparison of the two OMICS data sets

Since we acquired proteomic as well as transcriptomic data from 24-week-old db/db and wild-type mice in an analogous manner, we were interested in investigating how well changes in transcript translated to changes in protein. Briefly, FPKM or normalized abundance values were log-transformed, before a t-test between the respective glial and flow through values was calculated. Only genes/proteins with p<0.05 and a glia:flow through ratio of twofold were considered to be glia-specific. The final comparison encompassed genes/proteins that were glia-specific in at least one of the two datasets and detected in both. Further, we calculated the ratios between the respective wild-type and mutant 6-month-old animals for each dataset enabling us to directly compare the inter-genotype differences between datasets. Finally, we calculated a spearman correlation coefficient. This analysis and the corresponding scatter plots were done using the R programming language (R Core Team 2014).

### Electroretinogram recordings (ERGs)

Mice were dark adapted overnight and the ERG measurements performed in the morning under dim red light illumination. The mice were anesthetized with 100 mg/kg ketamine and 5 mg/kg xylazine by i.p. injection. The pupils of the mice were fully dilated with 0,5% tropicamide-phenylephrine 2,5% eye drops. Body temperature was maintained at 37 °C using a heating pad. Espion ERG Diagnosys equipment was used for the simultaneously recordings of both eyes. Corneal hydration was ensured by application of Methocel® 2% eyedrops. A reference and a ground needle electrode were positioned subcutaneously between the eyes and in the back, respectively. Golden loop electrodes were placed on the cornea. Rod-driven responses were measured and quantified at scotopic light conditions (0,001 cd/ms). Furthermore, mixed (rod- and cone-driven) responses were analyzed by applying light flashes of 3 cd/ms. Whereas cone-driven light responses (30 cd/ms) were recorded after 5 min of light adaption. Analysis of the data was done by using the Espion V6 software. A-wave amplitude was measured from baseline to the trough of the a-wave. B-wave amplitude was calculated from the trough of the a-wave to the peak of the b-wave.

### Intravitreal AAV administration

12-week-old mice were anesthetized with 100 mg/kg ketamine and 5 mg/kg xylazine by intraperitoneal injection. After the pupils of the mice were fully dilated with 0,5% tropicamide-phenylephrine 2,5% eye drops, 2 µl of AAV9-GFAP(0.7)-mNR3C1-2A-EGFP particles (5 × 10^13^ GC/ml; Vector Biolabs, AAV-266053) or PBS (sham control) were administered using the NanoFil™ injection system from World Precision Instruments (WPI). The AAV9-GFAP(0.7)-mNR3C1-2A-EGFP vector enables mNR3C1 and EGFP expression driven by the GFAP promoter with a 2A linker in between for co-expression. Methocel® 2% eyedrops were applied to the eyes of the mice after injection. 12 weeks post injection, morphometric analysis and ERG measurements were performed.

### Statistical analysis

The data were analyzed with GraphPad PRISM 8 and reported as mean ± standard error (SEM). Identification of outliers was also performed with GraphPad PRISM 8.

## Acknowledgements

We thank Dirkje Felder and Gabriele Jäger for excellent technical assistance. This research was supported by funding via the German Research Foundation (GR4403/2-1) and the ProRetina Foundation Germany (Pro-Re/Seed/Grosche.1-2014).

## Authors’ contributions

AMP generated, analyzed and interpreted data from phenotypic characterization, in vitro cortisol experiment, in vivo application of AAV-vectored GR overexpression and was a major contributor to the drafting of the manuscript. LK analyzed and interpreted the RNA-seq and MS/MS mass spectrometry data. MC generated the ATAC-seq dataset. FG analyzed the raw RNA-seq data. NDL generated and analyzed the trypsinized retinal flatmounts for the vascular phenotype of the mice. FG performed and analyzed the in vitro retinal explant culture experiment. KAW performed the super-resolution microscopy (STED) analysis to unambiguously localize GR in Müller cell nuclei. SG analyzed the microglial phenotype of AAV-transduced eyes. NK provided expertise, samples and mice for proper phenotyping of our db/db mouse strain. TP performed and analyzed patch clamp recordings on Müller cells from db/db mice. BHW interpreted the results of the study. SFK generated, analyzed, and interpreted the ERG recordings in this study. BB analyzed and interpreted the ATAC-seq dataset. SMH analyzed and interpreted the MS/MS mass spectrometry data. AG designed and supervised the study, analyzed and interpreted the data, and contributed substantially to the drafting of the manuscript. All authors read, revised, and approved the final manuscript.

## Availability of data and materials

All data supporting the results of this study are available in the article and its supplementary information. RNA-seq data have been deposited in the Gene Expression Omnibus (GEO) database under accession number GSE236627. The ATAC-seq data will also be made available via GEO. The mass spectrometry proteomics data have been deposited to the ProteomeXchange Consortium via the PRIDE (Perez-Riverol et al. 2022) partner repository with the dataset identifier PXD045085.

## Competing interests

The authors declare that they have no competing interests.

## Abbreviations

AAV: Adeno-associated virus
ATAC-seq: Assay for transposase-accessible chromatin using sequencing DR Diabetic retinopathy
EGFP: Enhanced green fluorescent protein ELISA Enzyme-linked immunosorbent assay ERG Electroretinogram
GCL: Ganglion cell layer
GFAP: Glial fibrillary acidic protein
GR: Glucocorticoid receptor
INL: Inner nuclear layer
IPL: Inner plexiform layer
MACS: Magnetic activated cell sorting ONL Outer nuclear layer
OPL: Outer plexiform layer OXPHOS Oxidative phosphorylation PBS Phosphate-buffered saline
qPCR: Quantitative realtime polymerase chain reaction TGFβ Tumor growth factor β
VEGF: Vascular endothelial growth factor

## Supplemental information

**Figure S1.**
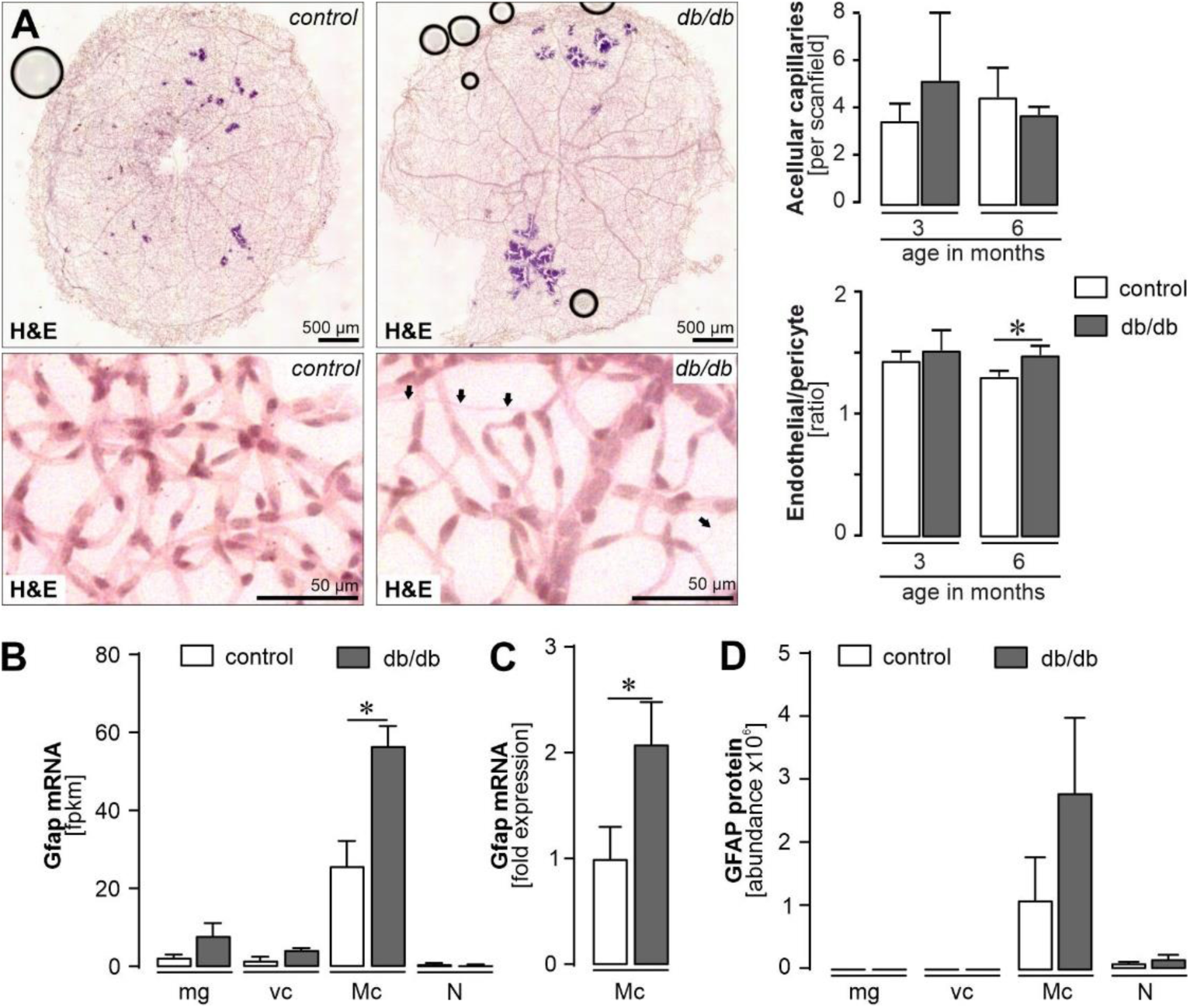
Validation of the vascular and glial alterations in the db/db mouse retina. *(A) Six-month-old db/db mice recapitulate some important vascular features of DR.* Left, *Representative images of hematoxylin and eosin (H&E) staining of trypsin-digested retinae from diabetic and control mice at 6 months of age. The arrows indicate acellular capillaries.* Right, *the number of acellular capillaries, endothelial cells, and pericytes per scan field was quantified in trypsin-digested flatmounts from db/db and control animals at 3 and 6 months of age. The ratio of endothelial cells to pericytes in the retina of db/db and control animals was then calculated using the respective cell counts. Bars represent mean ± SEM of results from n=3-4 animals per group. Unpaired t-test: *P<0.05. Scan field: 100.05 µm × 100.05 µm*. *(B) GFAP upregulation on transcript level in diabetic mice 6 months of age. RNA-seq was performed on RNA extracted from magnetic activated cell sorted (MACS) retinal cell types. Bars represent mean ± SEM (n=3-4). Unpaired t-test *P<0.05*. *(C) RNA-seq results were confirmed by qPCR on samples of MACS-sorted retinal cells isolated from 6-month-old mice*. *Bars represent mean ± SEM (n=5-7). Unpaired t-test: *P<0.05*. *(D) MS/MS mass spectrometry was used to assess the proteome of MACS-purified retinal cell types purified from retinae of 6-month-old mice. Bars represent mean ± SEM (n=4)*. *(B-D) Mg, microglia; vc, vascular cells; Mc, Müller cells; N, neurons*.

**Figure S2:**
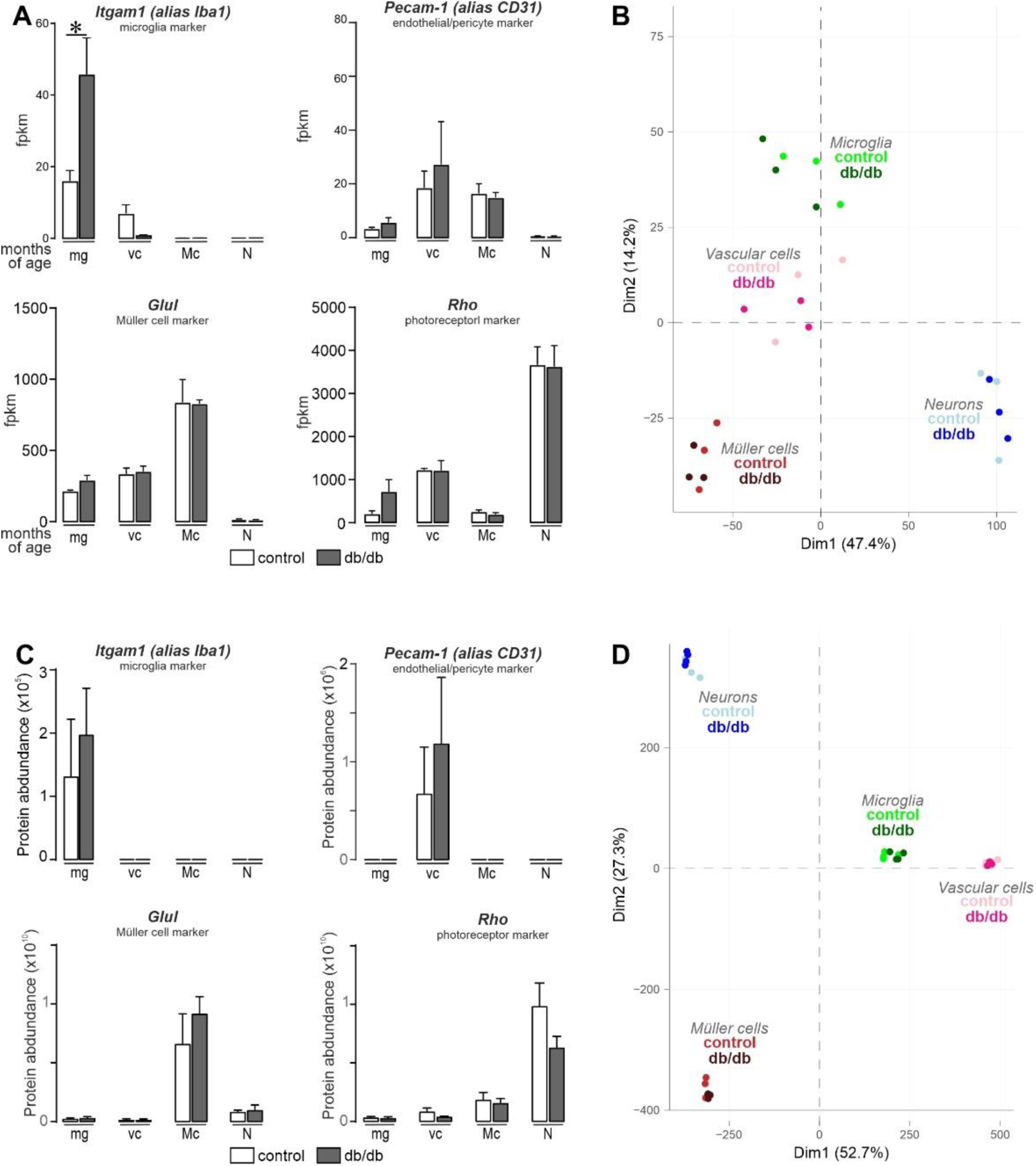
Validation of efficiency of MACS-based enrichment of retinal cell populations isolated from 6-month-old diabetic and control animals by RNA-seq data and tandem mass spectrometry. *(A) Plotting of mRNA expression levels of established cell markers as determined by RNA-seq demonstrates successful enrichment of retinal cell types from 6-month-old mice by magnetic activated cell sorting (MACS) irrespective of the genotype. Itgam, integrin subunit alpha M; Pecam1, platelet and endothelial cell adhesion molecule 1; Glul, glutamine synthetase; rho, rhodopsin. Bars represent mean ± SEM (cells purified from n=3-4 animals). Unpaired t-test: *P<0.05*. *(B) Principal component analysis (PCA) of the RNA-seq data from major retinal cell types enriched from retinae of control and diabetic mice at 6 months of age. Distinct clusters are formed by the different cell types implicating a good level of cell enrichment, while age and genotype seem to be less relevant for cluster formation*. *(C) The protein expression of marker genes for the four cell types implicated a successful separation of the different cell population from 6-month-old mice. MG: Microglia, VC: Vascular cells, MC: Müller cells; N: Neurons. GLUL: Glutamine synthetase. RHO, rhodopsin; ITGAM, integrin subunit alpha M; PECAM1, platelet and endothelial cell adhesion molecule 1. Bars represent mean ± SEM (n=4)*. *(D) Principal component analysis (PCA) of the proteomics data from four diabetic and control mice with an age of 6 month. Distinct clusters are formed by the four cell type groups*. *(A, C) mg, microglia; vc, vascular cells; Mc, Müller cells; n, neurons*.

**Figure s3.**
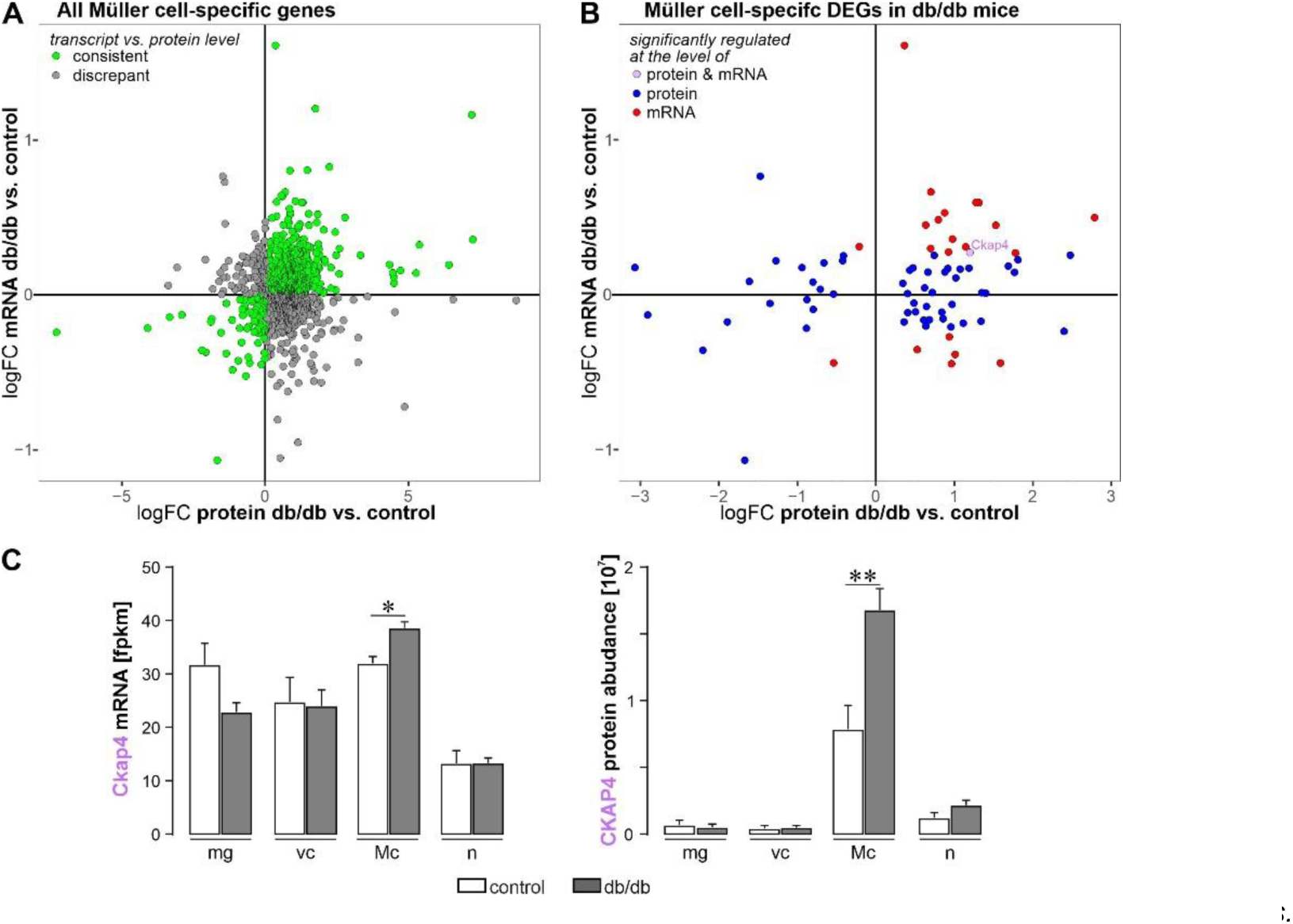
Comparison of Müller cell-specific genes in the diabetic retina at the transcriptome and proteome levels. *(A) Comparison of the results of RNA-seq and proteome analysis focusing on genes that are Müller cell-specific at either protein or mRNA level shows that out of the 883 genes consistently present in both data sets, 56.4% of the genes show a concordant expression pattern at the transcript and protein level*. *(B) Considering only the Müller cell-specific genes that additionally show significant differential expression in db/db mice at 6 months of age, 57.3% of the 75 genes show a concordant regulation pattern at transcript and protein level. Unpaired t-test: *P<0.05., **P<0.01*. *(C) MRNA and protein abundance of CKAP4 in microglia (mg), vascular cells (vc), Müller cells (Mc) and retinal neurons (n) as determined by RNA-seq and MS/MS mass spectrometry. Bars represent mean ± SEM (n=3-4). Unpaired t-test: *P<0.5; **O<0.01*.

**Figure s4.**
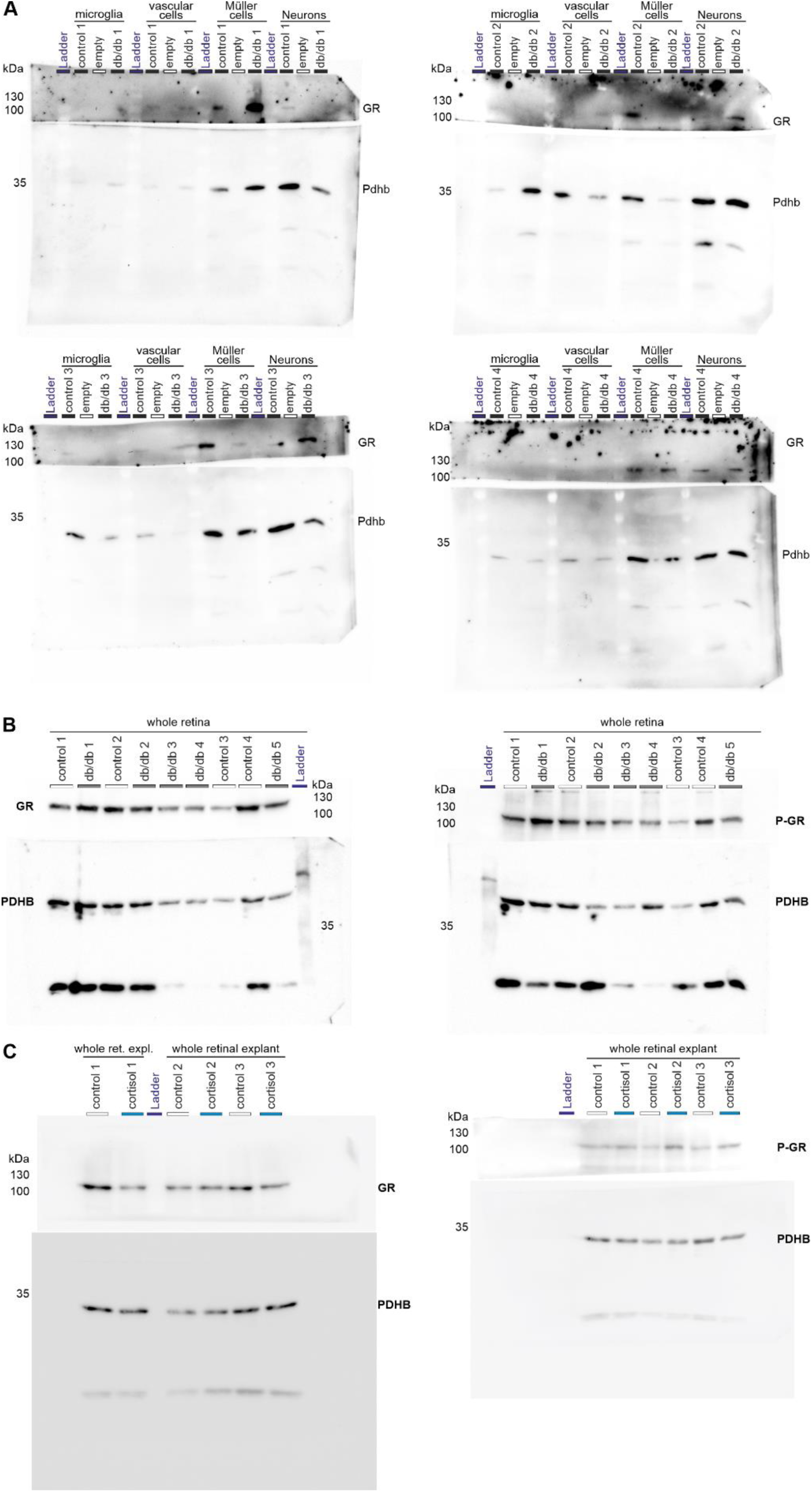
*Overview of all uncropped Western blots used for the analysis shown in* Figures 5 *and 6.* *(A) Western blots for the analysis shown in* Figure 5B*. Each blot was cut at the 70 kDa band of the ladder to detect the high molecular weight GR (95 kDa) and the low molecular weight housekeeper PDHB (∼34 kDa) on the same blot. Since the amount of protein per MACS-purified cell population was very low, the amount of protein loaded per lane was not adjusted (the whole extract per cell population was loaded), but normalized to the housekeeper expression*. *(B) Western blots for the analysis shown in* Figure 5D*. Each blot was cut at the 70 kDa band of the ladder to detect GR (95 kDa) or its phosphorylated form and the housekeeper PDHB (∼34 kDa) on the same blot. 50 µg of protein extract from whole retinae was loaded per lane and expression levels were normalized to that of the housekeeper*. *(C) Western blots for the analysis shown in* Figure 6B*. Each blot was cut at the 70 kDa band of the ladder to detect GR (95 kDa) or its phosphorylated form and the housekeeper PDHB (∼34 kDa) on the same blot. 50 µg of protein extract per retinal explant culture was loaded per lane and expression levels were normalized to that of the housekeeper*.

## Notes

### Competing Interest Statement

The authors have declared no competing interest.

### Summary of Updates

Affiliation of authors corrected.

